# Candidate Molecular Subtypes of Cognitive Resilience in Alzheimer’s Disease: A Multi-Cohort Machine Learning and Neuroimaging Study

**DOI:** 10.1101/2024.08.25.609610

**Authors:** Akihiro Kitani, Yusuke Matsui, the Alzheimer’s Disease Neuroimaging Initiative

**Affiliations:** Institute for Glyco-core Research (iGCORE), Nagoya University, 461-8673 Nagoya, Aichi, Japan; Biomedical and Health Informatics Unit, Department of Integrated Health Science, Nagoya University Graduate School of Medicine, Nagoya, Japan

**Keywords:** Alzheimer’s disease, cognitive resilience, molecular subtypes, transcriptomics, proteomics, machine learning, neuroimaging

## Abstract

**Background:** Cognitive resilience (CR) in Alzheimer’s disease (AD) refers to preserved cognitive function despite substantial AD pathology. Diverse biological processes have been implicated in CR, including synaptic maintenance, neuroimmune regulation, and metabolic homeostasis. However, how these mechanisms are organized into molecularly distinct CR subtypes and relate to clinical and neuroanatomical heterogeneity remains unclear. Here, we applied a machine learning framework to multi-cohort transcriptomic, proteomic, and neuroimaging data to investigate molecular subtypes of CR in AD.

**Methods:** RNA-seq data from the Religious Orders Study and Memory and Aging Project (ROSMAP) cohort were used to train machine learning models classifying individuals with AD pathology as CR or non-CR based on residual-based resilience scores. Model development and performance estimation used nested cross-validation to minimize information leakage. Final ROSMAP-trained models were evaluated in the independent Mount Sinai Brain Bank (MSBB) cohort. Model-derived genes were used for biological interpretation and hierarchical clustering of CR individuals. The subtype structure was further evaluated in the Alzheimer’s Disease Neuroimaging Initiative (ADNI) cohort using cerebrospinal fluid proteomics, MRI-derived brain measures, and longitudinal MMSE data.

**Results:** Machine learning models showed modest but consistent predictive performance in ROSMAP, with out-of-fold AUROC values of 0.644–0.688. In the independent MSBB full cohort, AUROC values were 0.586–0.659, with improved discrimination in a top/bottom quartile analysis. Hierarchical clustering identified two major molecular subgroups among CR individuals in ROSMAP/MSBB RNA-seq data. A reduced 22-gene/protein signature showed a partial, cluster-like resemblance to this structure in ADNI cerebrospinal fluid proteomics. In ADNI, both projected CR subtypes showed preserved brain tissue-volume profiles and slower longitudinal MMSE decline compared with non-CR participants, whereas clear differences between CR subtypes were not observed. Differential CSF proteomic analysis suggested partially distinct molecular characteristics.

**Conclusions:** These findings suggest that CR in AD may encompass molecularly heterogeneous, subtype-like profiles that converge on broadly preserved brain structure and slower cognitive decline. Our results provide a candidate framework for stratifying resilience-associated molecular phenotypes in AD and warrant prospective and experimental validation. We also developed the Resilience Gene Analyzer, a web-based platform for visualizing gene-level contributions to CR prediction (https://igcore.cloud/GerOmics/REsilienceGeneAnalyzer/).

## Background

Alzheimer’s disease (AD) remains a formidable challenge due to its multifactorial and incompletely understood etiology, combined with the urgent need for effective interventions. In AD, cognitive resilience (CR) refers to the capacity of some individuals to sustain cognitive performance despite the accumulation of hallmark AD pathologies—amyloid plaques and neurofibrillary tangles. CR is typically quantified as the deviation between observed cognitive function and the level predicted from neuropathological burden [1–3]. Multiple demographic, anatomical, and psychosocial factors — including sex/gender differences [4]; educational attainment, and brain weight [5–7]; personality traits; comorbid Parkinson’s disease and depression; life activities; and eudaimonic well-being— have been implicated [8–10]. However, the molecular mechanisms underlying CR remain largely unresolved.

Recent studies have begun to elucidate the molecular mechanisms underlying CR. High-throughput omics technologies have transformed AD research by enabling comprehensive interrogation of disease mechanisms, biomarker discovery, and drug target identification [11–14]. Genomic studies have associated CR with variants in genes such as APOE, BDNF, and Klotho [15–22], while transcriptomic and proteomic studies have implicated proteins including NRN1, ACTN4, YWHAG, and NPTX2 [23–29]. Emerging evidence suggests that CR may be supported by diverse biological processes, including synaptic preservation and maintenance of neuronal activity and functional connectivity, suppression of excessive neuroinflammation, and preservation of mitochondrial and metabolic homeostasis. However, these mechanisms are unlikely to be uniformly shared among all resilient individuals. Instead, CR may arise through multiple, partially independent molecular pathways that converge on preserved cognition despite AD pathology. Despite these advances, whether such heterogeneity can be organized into molecularly distinct CR subtypes—and how these subtypes relate to clinical and neuroanatomical features—remains unclear. Addressing this knowledge gap is essential for understanding the biological diversity of CR and for developing candidate stratification frameworks for future mechanistic and translational studies [1,30,31].

Machine learning (ML) has emerged as a powerful framework for extracting clinically relevant signatures from high-dimensional omics data, enabling disease prediction, patient stratification, and mechanistic insight [32–36]. While ML approaches have been successfully applied to AD diagnosis and prognosis, their potential to dissect the molecular architecture of cognitive resilience has remained underexplored.

Here, we integrate ML with multi-cohort transcriptomic, proteomic, and neuroimaging data analysis to investigate and characterize molecular subtypes of CR in AD. Using RNA-seq data from the Religious Orders Study and Memory and Aging Project (ROSMAP) [37], we developed predictive models of CR status and evaluated their cross-cohort generalizability in the independent Mount Sinai Brain Bank (MSBB) dataset [38]. To delineate CR heterogeneity, we performed feature-based clustering and identified two molecular subgroups among CR individuals. We then evaluated the subtype structure in the Alzheimer’s Disease Neuroimaging Initiative (ADNI) cohort using cerebrospinal fluid (CSF) proteomics. In ADNI, both projected CR subtypes showed preserved global brain tissue-volume profiles and slower longitudinal cognitive decline compared with non-CR individuals. Finally, we developed the Resilience Gene Analyzer (REGA), a publicly accessible platform for visualizing model-derived gene-level contribution scores for CR prediction, including ElasticNet coefficients and SHAP-based attributions from non-linear models [39]. These results advance the understanding of molecular heterogeneity in CR and may inform future personalized approaches in AD.

## Results

### Cross-cohort Evaluation of Cognitive Resilience Prediction Models

An overview of the analytic pipeline is illustrated in Fig. 1. To investigate molecular subtypes of cognitive resilience (CR) in Alzheimer’s disease (AD), we first developed machine learning (ML) models to classify individuals as CR or non-CR (Fig. 1a). Feature analysis was then applied to define candidate molecular subtype signatures among CR individuals. We subsequently examined whether analogous subtype-like profiles could be detected in the Alzheimer’s Disease Neuroimaging Initiative (ADNI) cohort by projecting the ROSMAP/MSBB-derived signature onto cerebrospinal fluid (CSF) proteomics data, and evaluated the resulting groups in relation to demographics and MRI-derived metrics (Fig. 1b).

**Fig. 1.**
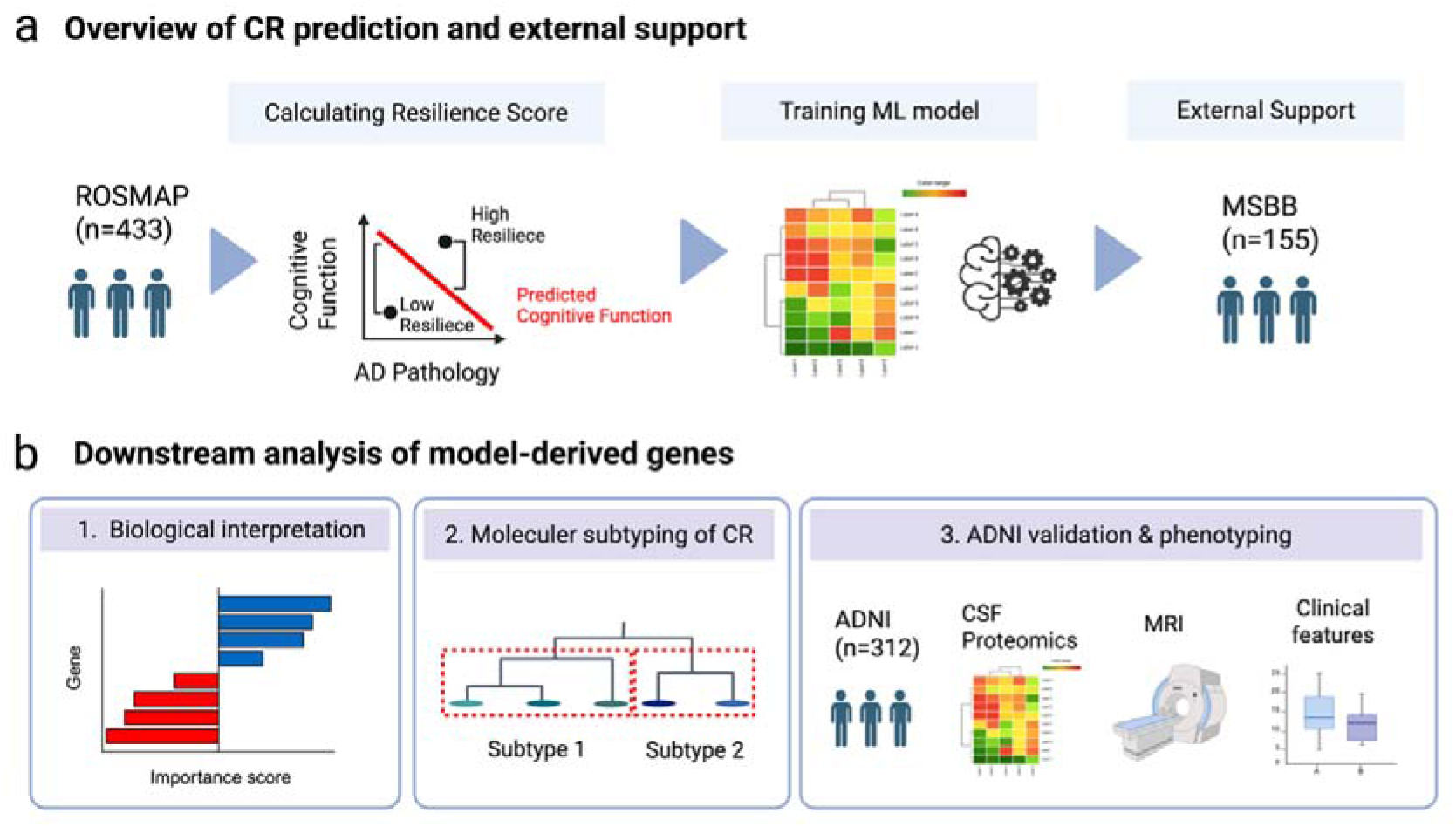
Overview of this study. (a) Machine learning models were constructed to classify CR and non-CR cases using RNA-seq data, with the ROSMAP [37] cohort, and their cross-cohort performance was evaluated in the independent MSBB [38] cohort. (b) Feature genes identified by the models were used for (1) biological interpretation to elucidate molecular mechanisms and (2) hierarchical clustering to investigate subtypes. (3) The subtype signatures were projected onto ADNI CSF proteomics to assess analogous subtype-like profiles.

In line with the conventional definition of CR as preserved cognitive function despite AD pathology, we restricted analyses to individuals with evidence of AD neuropathology in both cohorts (n = 433 in ROSMAP and n = 155 in MSBB; see Methods for details). Residual-based resilience scores were calculated from the relationship between cognitive performance and neuropathological burden, using MMSE in ROSMAP and CDR in MSBB as cohort-specific cognitive measures. Including age, sex, and years of education as additional covariates had minimal impact on the resulting resilience scores (Supplementary Fig. 1; see Methods). To improve cross-cohort model generalizability, batch effects in gene expression were corrected across datasets (Supplementary Figs. 2,3). Post-correction principal component analysis (PCA) suggested that major variation in the corrected expression data was not visually dominated by cohort, APOE genotype, age, or sex in the leading principal components (Supplementary Fig. 3c–f).

Using the batch-corrected gene expression data, we developed four ML classifiers—ElasticNet [40], support vector machine (SVM) [41], Random Forest [42], and extreme gradient boosting(XGBoost) [43] on the ROSMAP dataset to classify individuals as CR or non-CR. To minimize information leakage, model development and primary performance estimation were performed in ROSMAP using nested cross-validation; feature selection, hyperparameter tuning, and feature-number selection were all confined to the inner cross-validation loop of each outer training set (Supplementary Fig. 4; see Methods for details).

Using out-of-fold predictions from ROSMAP nested cross-validation, the models showed modest but consistent predictive performance, with AUROC values of 0.679, 0.688, 0.677, and 0.644 for ElasticNet, SVM, Random Forest, and XGBoost, respectively. Corresponding AUPRC values were 0.728, 0.735, 0.729, and 0.711 (Fig. 2a, Supplementary Fig. 5a, Supplementary Table S1). Additional metrics, bootstrap 95% confidence intervals, Brier scores, and calibration curves are shown in Fig. 2 and Supplementary Fig. 5. Selected model configurations are summarized in Supplementary Table S2.

**Fig. 2.**
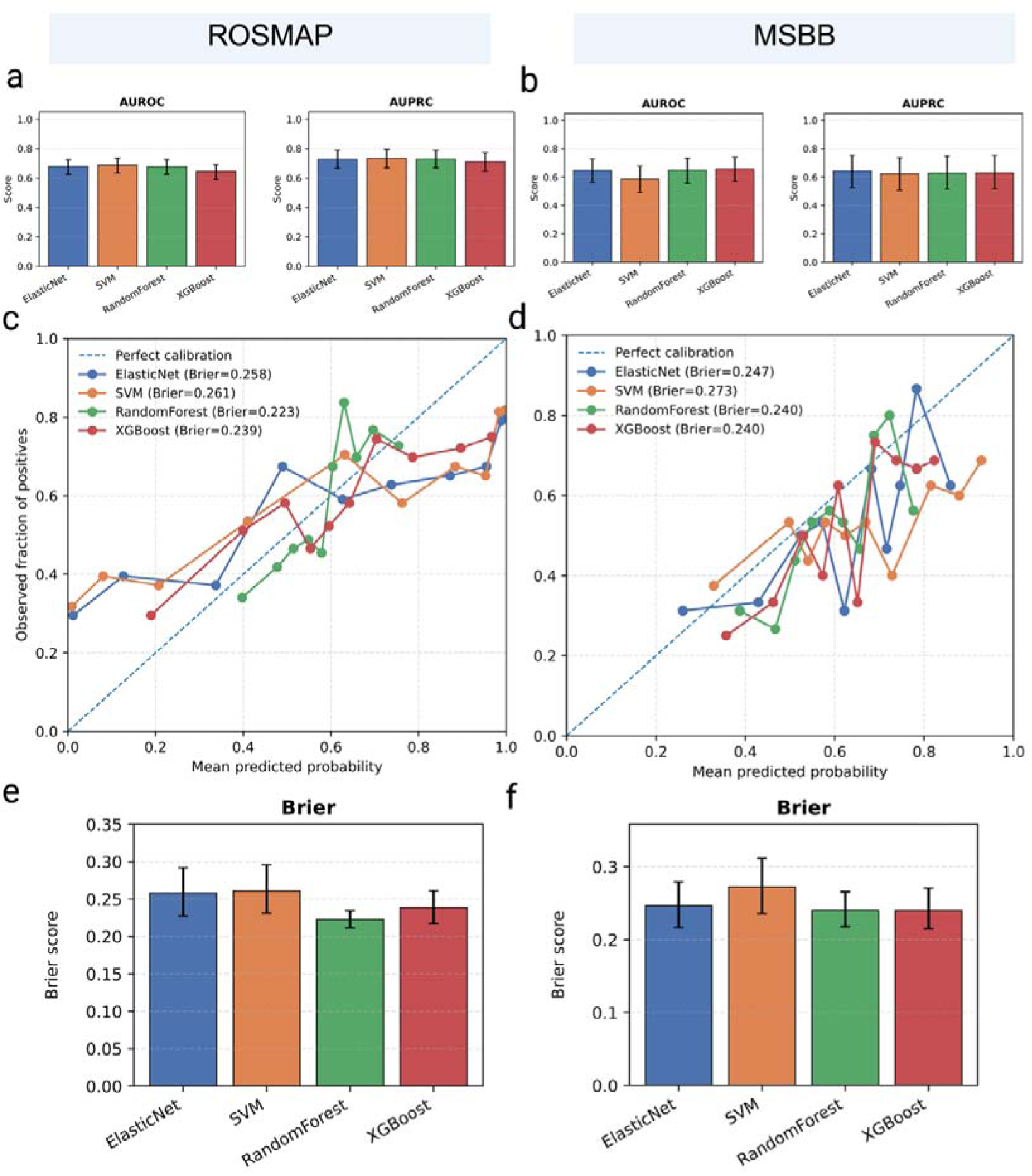
Predictive performance and calibration of CR classification models in ROSMAP and MSBB. (a, b) AUROC and AUPRC of four machine learning models—ElasticNet, support vector machine (SVM), Random Forest, and XGBoost—for classifying individuals as cognitive resilience (CR) or non-CR. Model performance was evaluated using out-of-fold predictions from nested cross-validation in ROSMAP (a) and independent evaluation in the full MSBB cohort (b). Bars indicate point estimates, and error bars indicate bootstrap 95% confidence intervals. (c, d) Calibration curves for the four models in ROSMAP nested cross-validation (c) and MSBB full external validation (d). The dashed line represents perfect calibration. Brier scores are shown in parentheses; lower values indicate better calibrated probability predictions. (e, f) Brier scores for the four models in ROSMAP nested cross-validation (e) and MSBB full external validation (f). Bars indicate point estimates, and error bars indicate bootstrap 95% confidence intervals.

Final models were then trained on the full ROSMAP cohort and applied once to the independent MSBB full cohort. MSBB was not used for feature selection, hyperparameter tuning, feature-number selection, or model selection (Supplementary Fig. 4). In the MSBB full cohort, AUROC values were 0.647, 0.586, 0.649, and 0.659, and AUPRC values were 0.643, 0.624, 0.629, and 0.630 for ElasticNet, SVM, Random Forest, and XGBoost, respectively (Fig. 2b, Supplementary Fig. 5b, Supplementary Table S1). This attenuation relative to ROSMAP may partly reflect cross-cohort differences in CR phenotype definition, including the use of MMSE in ROSMAP and CDR in MSBB.

As a secondary sensitivity analysis, we evaluated the same fixed models in the MSBB top/bottom quartile subset, where AUROC values increased to 0.775, 0.641, 0.756, and 0.751 for ElasticNet, SVM, Random Forest, and XGBoost, respectively (Supplementary Fig. 5c, Supplementary Table S1). Although this quartile-based analysis was not used for model selection, the improved discrimination in individuals with more pronounced CR phenotypes suggests that the models captured CR-related transcriptomic signals.

### Model-derived Key Features Are Linked to AD-related Biological Processes

Given the modest but consistent predictive performance of the ML models, we next explored whether the model-derived features showed biologically interpretable patterns, without assuming that they directly represent causal CR mechanisms. Features from each model were categorized as positive or negative contributors to CR prediction based on model-specific contribution scores, including ElasticNet coefficients and SHAP-based feature attributions for the other models (see Methods). After combining non-redundant features across the four models, we identified 371 positively contributing genes and 352 negatively contributing genes (Supplementary Fig. 6; Supplementary Table S3).

Several of the top-ranked model-derived genes have been previously implicated in AD-related biological processes, including oxidative stress, synaptic and axonal function, mitochondrial regulation, neuroinflammation, and cell-death pathways. For example, ALDH1A1 [44] and NOX4 [45] have been linked to oxidative stress responses and Aβ-associated toxicity in AD-related contexts, while KIF5A and KIF5C are involved in kinesin-mediated axonal transport, a process closely related to synaptic maintenance and tau-associated neuronal dysfunction [46,47]. SRR, which regulates D-serine/NMDA receptor signaling, and PDE10A, which is connected to cAMP–PKA–CREB signaling, may reflect pathways relevant to synaptic plasticity and neuronal resilience [48,49]. In contrast, genes such as CASP8 may represent inflammatory or cell-death-related mechanisms rather than protective resilience mechanisms per se [50]. These observations suggest that, although the model-derived genes should not be interpreted as causal mediators of CR without further validation, the highest-ranking features include genes with plausible links to AD pathophysiology and resilience-related biological processes.

### Hierarchical Clustering Suggested Candidate Molecular Heterogeneity among CR individuals

To investigate molecular heterogeneity among individuals with cognitive resilience (CR), we performed hierarchical clustering of CR samples from the ROSMAP and MSBB cohorts using 144 model-derived genes with positive contributions to CR prediction. The 144-gene expression profile separated CR individuals into two major subgroups (Fig. 3a). To facilitate cross-platform comparison with ADNI CSF proteomics, we focused on the 22 genes that overlapped with proteins measured in the ADNI SOMAscan dataset. These shared genes retained subtype-associated expression differences in the ROSMAP/MSBB RNA-seq data (Fig. 3b; Supplementary Table S4), supporting their use as a reduced signature for subsequent projection into ADNI. For example, KL [22], which encodes the CR-associated protein Klotho, and PDGFD [51] showed higher expression in cluster 2, whereas SULT1A1, which has recently been implicated in AD-related phenotypes [52] and cognitive function [53], showed higher expression in cluster 1 (Fig. 3b).

**Fig. 3.**
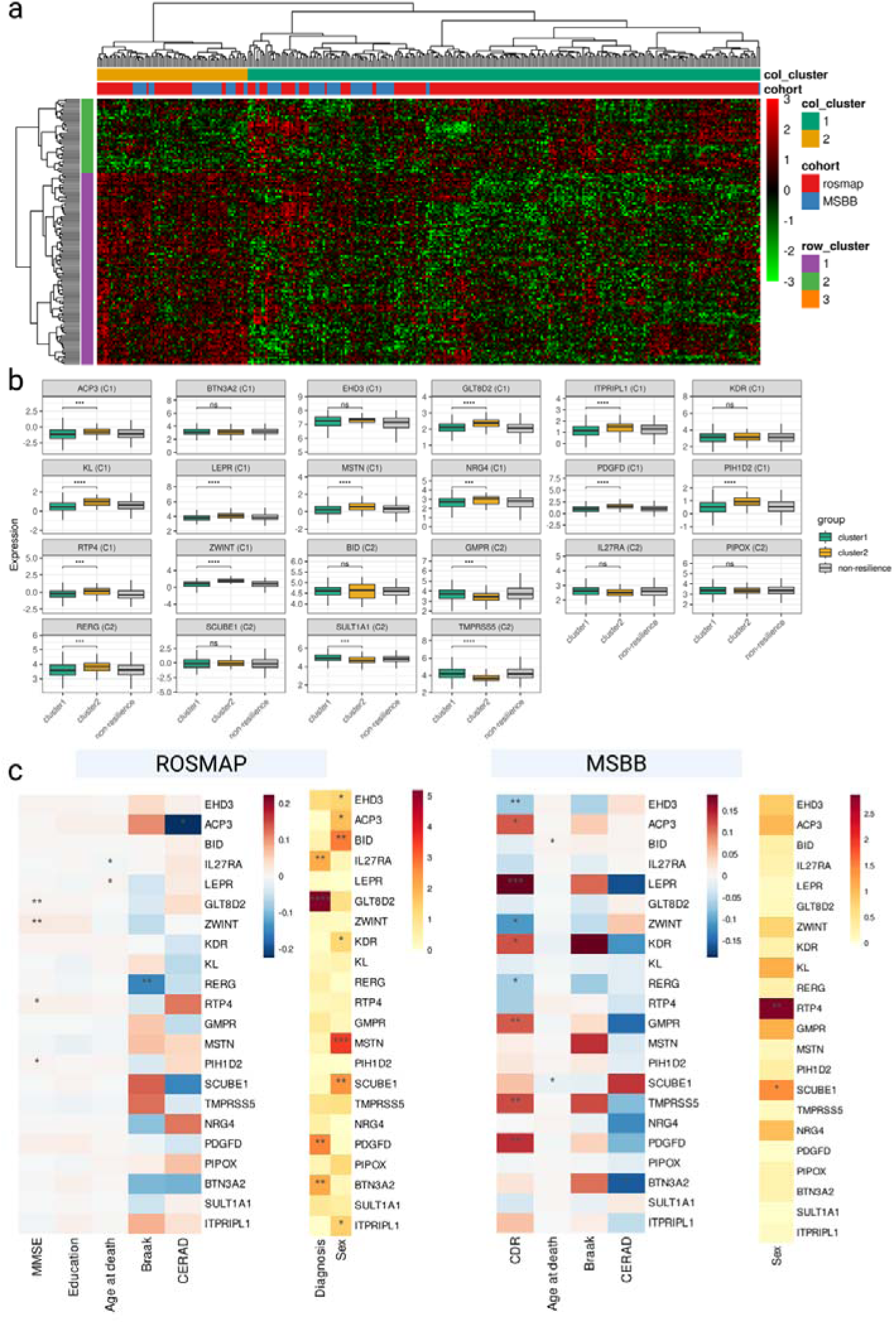
Hierarchical clustering identified molecular subtypes among CR individuals in ROSMAP and MSBB RNA-seq data. (a) Heatmap of hierarchical clustering; rows represent genes and columns represent samples. (b) Representative gene expression levels among CR cluster 1, cluster 2, and non-CR groups. Statistical significance between cluster 1 and cluster 2 was assessed by t-test with FDR correction. (c) Linear-model associations between subtype-associated gene expression and clinical or neuropathological variables in the ROSMAP and MSBB cohorts. Heatmap colors indicate regression coefficients for continuous variables or −log10 raw P values for categorical variables. (*: p□<□0.05, **: p□<□0.01, ***: p□<□0.001, ****: p□<□0.0001)

We further examined the relationships between these subtype-associated genes and clinical or neuropathological traits. In the ROSMAP and MSBB cohorts, gene expression levels showed variable associations with cognitive performance, educational attainment, diagnosis, and neuropathological measures, including MMSE, years of education, CERAD score, Braak score, and CDR score, with both positive and negative directions depending on the gene and phenotype (Fig. 3c). These findings suggest that the two CR subgroups identified in the ROSMAP and MSBB cohorts may reflect distinct molecular programs associated with cognitive resilience and support molecular heterogeneity within CR.

To evaluate whether the subtype structure was partially reflected, we next analyzed cerebrospinal fluid (CSF) proteomics data from the ADNI cohort. Among the 144 genes used in the ROSMAP and MSBB cohort analysis, 22 genes had corresponding proteins measured in the ADNI SOMAscan dataset. Before projecting the ROSMAP/MSBB subtype signatures to ADNI, we confirmed that these 22 shared genes retained subtype-associated expression differences in the ROSMAP/MSBB RNA-seq data (Supplementary Table S4), supporting their use as a reduced cross-platform signature.

We next examined whether the 22 shared proteins in ADNI were associated with clinical and pathological phenotypes among ADNI individuals with AD pathology and high CR. Linear-model analyses showed that multiple proteins were associated with cognitive performance, resilience score, CSF AD biomarkers, diagnosis, and demographic variables, including MMSE, Aβ42, Tau, pTau, age, education, and sex (Fig. 4a).

**Fig. 4.**
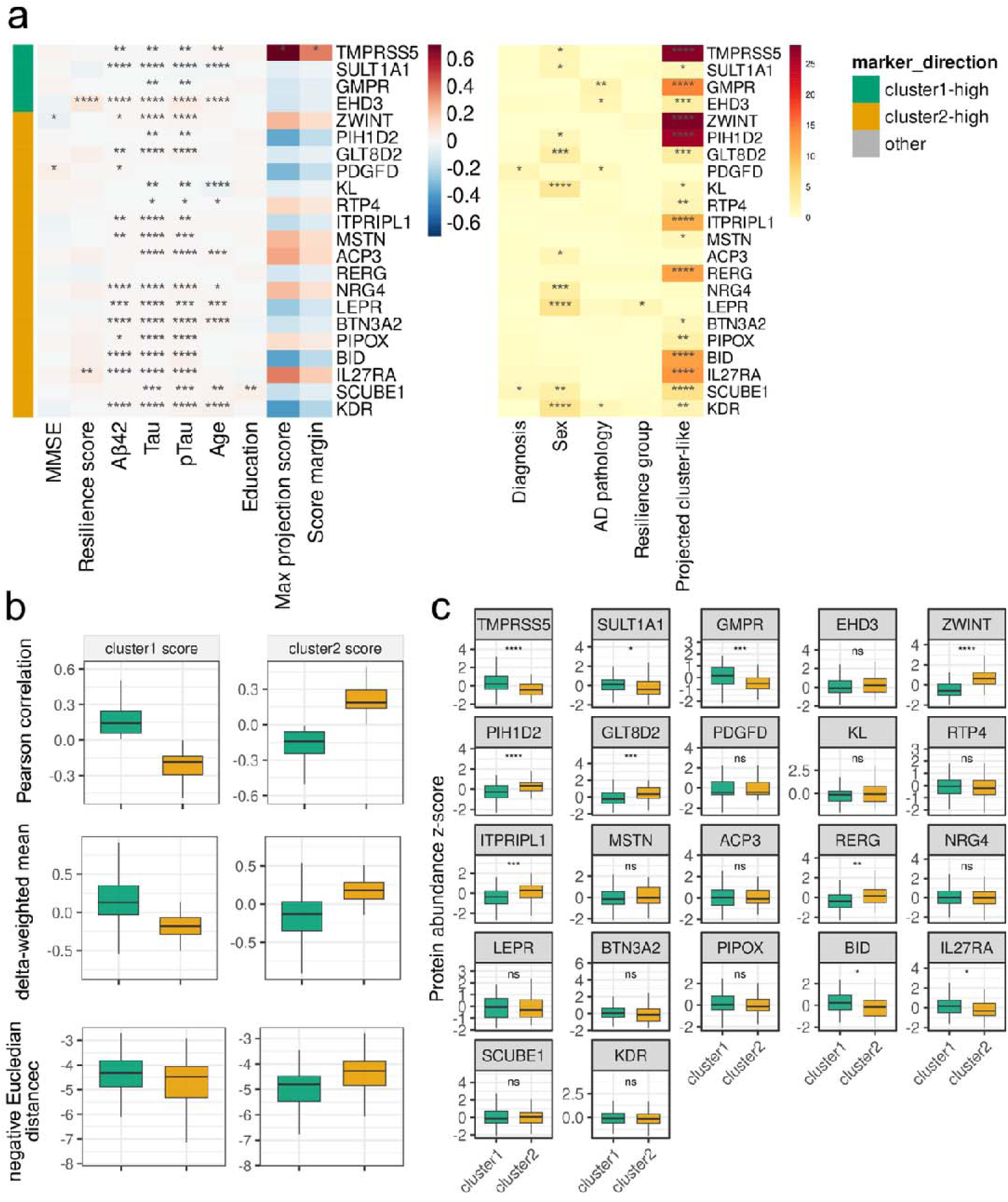
ADNI CSF proteomics provides cross-platform support for the ROSMAP/MSBB-derived CR subtype structure. (a) Linear-model associations between the 22 shared ADNI CSF proteins and clinical, demographic, and biomarker variables. (b) Projection score distributions calculated using centroid correlation, marker-delta–weighted scoring, and negative Euclidean distance. (c) Protein abundance differences between projected cluster1-like and cluster2-like ADNI samples for the 22 shared proteins. Statistical significance between cluster 1 and cluster 2 was assessed by t-test with FDR correction. (*: p□<□0.05, **: p□<□0.01, ***: p□<□0.001, ****: p□<□0.0001)

To evaluate whether the ROSMAP/MSBB-derived subtype structure was reflected in ADNI, we calculated subtype projection scores for each ADNI individual. Briefly, cluster-specific centroids were computed from the ROSMAP/MSBB expression profiles of the 22 shared genes, and each ADNI CSF proteomic profile was compared with these centroids. ADNI samples were assigned as cluster1-like or cluster2-like according to the centroid with the higher similarity score (see Methods for details). Using this centroid-based projection, ADNI individuals were assigned to cluster1-like or cluster2-like groups. The assigned groups showed distinct centroid-correlation scores, and similar separation was observed using alternative scoring approaches, including marker-delta–weighted scoring and negative Euclidean distance (Fig. 4b), indicating that the projection was not solely dependent on a single scoring metric.

Finally, we examined whether individual proteins showed expression differences between the projected ADNI cluster-like groups. Several of the 22 shared proteins showed directionally consistent differences between cluster1-like and cluster2-like groups, including proteins corresponding to ROSMAP/MSBB subtype marker directions (Fig. 4c; Supplementary Table S5). Although the ADNI analysis was based on CSF proteomics rather than brain RNA-seq and was restricted to a reduced 22-protein overlap, these results provide partial but consistent cross-cohort and cross-platform support for the ROSMAP/MSBB-derived CR subtype structure. Additionally, no significant differences in age, sex distribution, or APOE ε4 carrier status were observed between the projected CR clusters, although years of education differed between cluster 2 and the non-CR group (Supplementary Fig. 7). These results suggest that the projected CR clusters were not primarily explained by major demographic variables.

### Structural and Longitudinal Cognitive Profiles of ADNI-projected CR Subtypes

To characterize the structural and longitudinal cognitive profiles of the ADNI-projected CR subtypes, we compared MRI-derived UCD (University of California, Davis) tissue-volume and WMH measures [60–63] among cluster 1, cluster 2, and non-CR participants in the ADNI cohort. The ADNI analysis subsets used for CSF proteomics, MRI-derived imaging measures, and longitudinal MMSE analyses included the same projected group sizes: cluster 1, n = 80; cluster 2, n = 70; and non-CR, n = 162. In covariate-adjusted linear models, both CR clusters showed evidence of preserved global tissue-volume profiles relative to the non-CR group (Fig. 5a; Supplementary Table S6). Specifically, compared with non-CR participants, both cluster 1 and cluster 2 showed lower CSF volume and higher gray matter and cerebrum brain volume measures after adjustment for age at MRI, sex, years of education, APOE ε4 carrier status, ADNI phase, scanner manufacturer, magnetic field strength when estimable, and total cerebrum cranial volume as an intracranial-size proxy. In contrast, hippocampal volume and WMH burden did not remain significant after multiple-testing correction. Importantly, none of the UCD imaging metrics showed a significant difference between cluster 1 and cluster 2 after FDR correction, suggesting that broad volumetric preservation was shared across the two CR subtypes rather than being specific to one subtype.

**Fig. 5.**
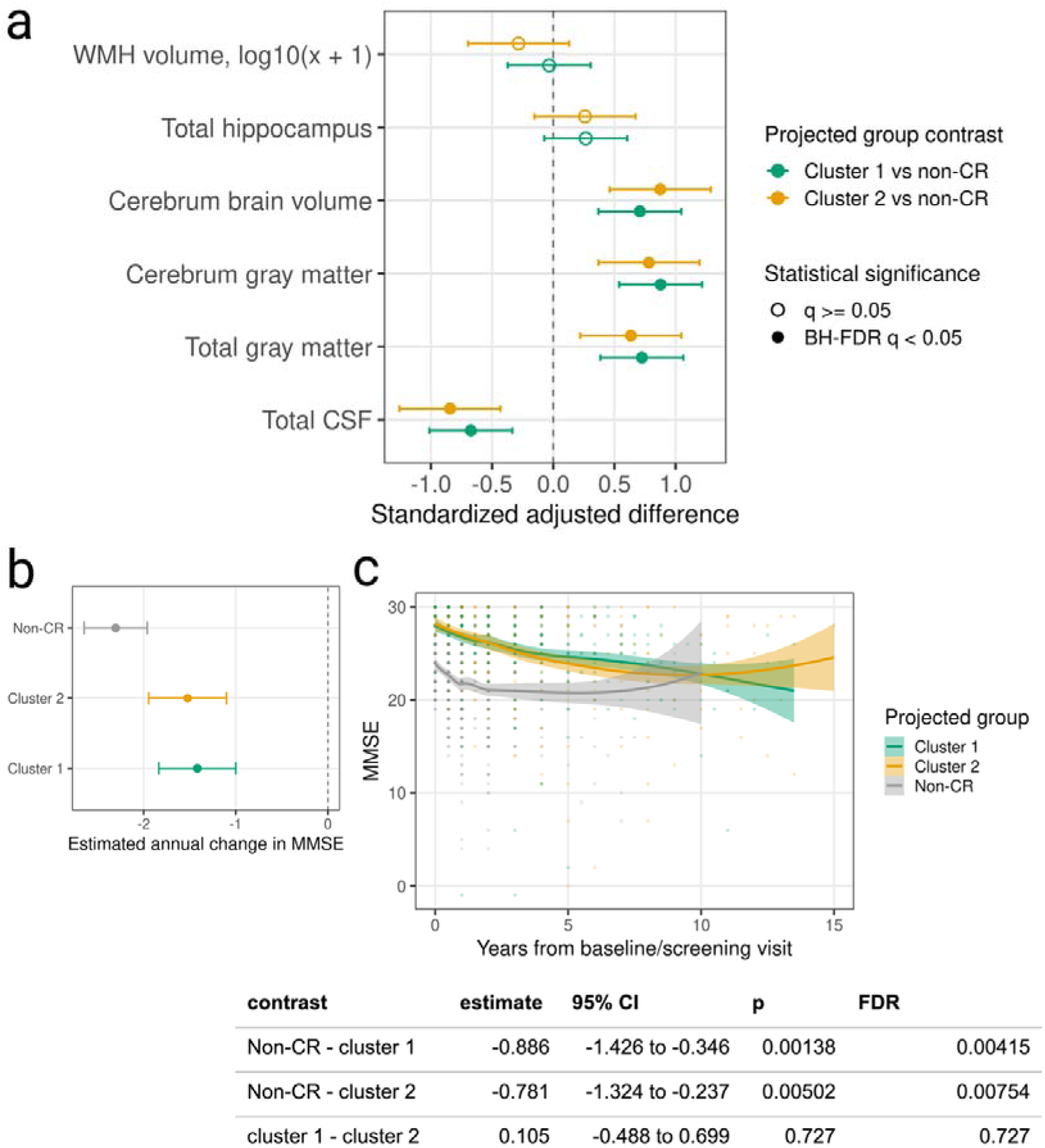
Structural and longitudinal cognitive profiles of ADNI-projected CR subtypes. (a) Forest plot of covariate-adjusted differences in UCD WMH and global tissue-volume measures for cluster 1 and cluster 2 relative to the non-CR group. Points indicate standardized adjusted differences, and horizontal bars indicate 95% confidence intervals. Filled points indicate BH-FDR q < 0.05. (b) Estimated annual MMSE change derived from linear mixed-effects models. Points indicate estimated slopes, and horizontal bars indicate 95% confidence intervals. (c) Longitudinal MMSE trajectories by projected group. Lines and shaded areas show model-fitted trajectories and 95% confidence intervals for visualization. The table summarizes pairwise contrasts in annual MMSE slopes.

As a complementary regional analysis, UCSF (University of California, San Francisco) FreeSurfer metrics [54–60] showed a broadly similar pattern of structural preservation in the CR clusters relative to the non-CR group (Supplementary Table S7).

We next evaluated whether the two CR clusters differed in longitudinal cognitive trajectories. Linear mixed-effects models of repeated MMSE measurements showed that the non-CR group had significantly steeper annual MMSE decline than either CR cluster (Fig. 5b,c; Supplementary Table S8). The estimated annual slope difference was −0.886 points/year for non-CR versus cluster 1 and −0.781 points/year for non-CR versus cluster 2. In contrast, annual MMSE slopes did not differ significantly between cluster 1 and cluster 2, indicating that the two molecularly defined CR subtypes were not primarily distinguished by differences in longitudinal cognitive decline rates. Cox proportional hazards models adjusted for demographic covariates were provided as supplementary time-to-decline analyses and showed a consistent pattern (Supplementary Table S9). Overall, these findings indicate that both CR subtypes show preserved global brain tissue measures and relatively slower cognitive decline compared with non-CR participants, whereas clear structural or longitudinal cognitive differences between cluster 1 and cluster 2 were not observed.

### Partially Distinct Molecular Characteristics of CR Subtypes

To further characterize the molecular features of the projected CR subtypes, we compared CSF proteomics between each CR subtype and the non-CR group (Fig. 6a). In cluster 2, NPTX2 levels were significantly higher than in the non-CR group, consistent with the prior report [29]. In contrast, cluster 1 showed a larger set of proteins with significantly lower expression than the non-CR group, whereas such decreases were largely absent in cluster 2. The sets of proteins differentially expressed in clusters 1 and 2 showed minimal overlap (Fig. 6b). Enrichment analysis of proteins with higher expression revealed predominant enrichment of neuronal pathways such as synapse assembly in both clusters (Fig. 6c, d; Supplementary Table S10). Proteins with lower expression in cluster 1 were largely enriched in immune-related pathways, including inflammatory response and phagocytosis (Fig. 6e). Collectively, these findings suggest that cluster 1 may be characterized by preserved neuronal and synaptic functions accompanied by reduced neuroinflammatory activity, whereas cluster 2 may be characterized primarily by enrichment of neuronal and synaptic maintenance pathways.

**Fig. 6.**
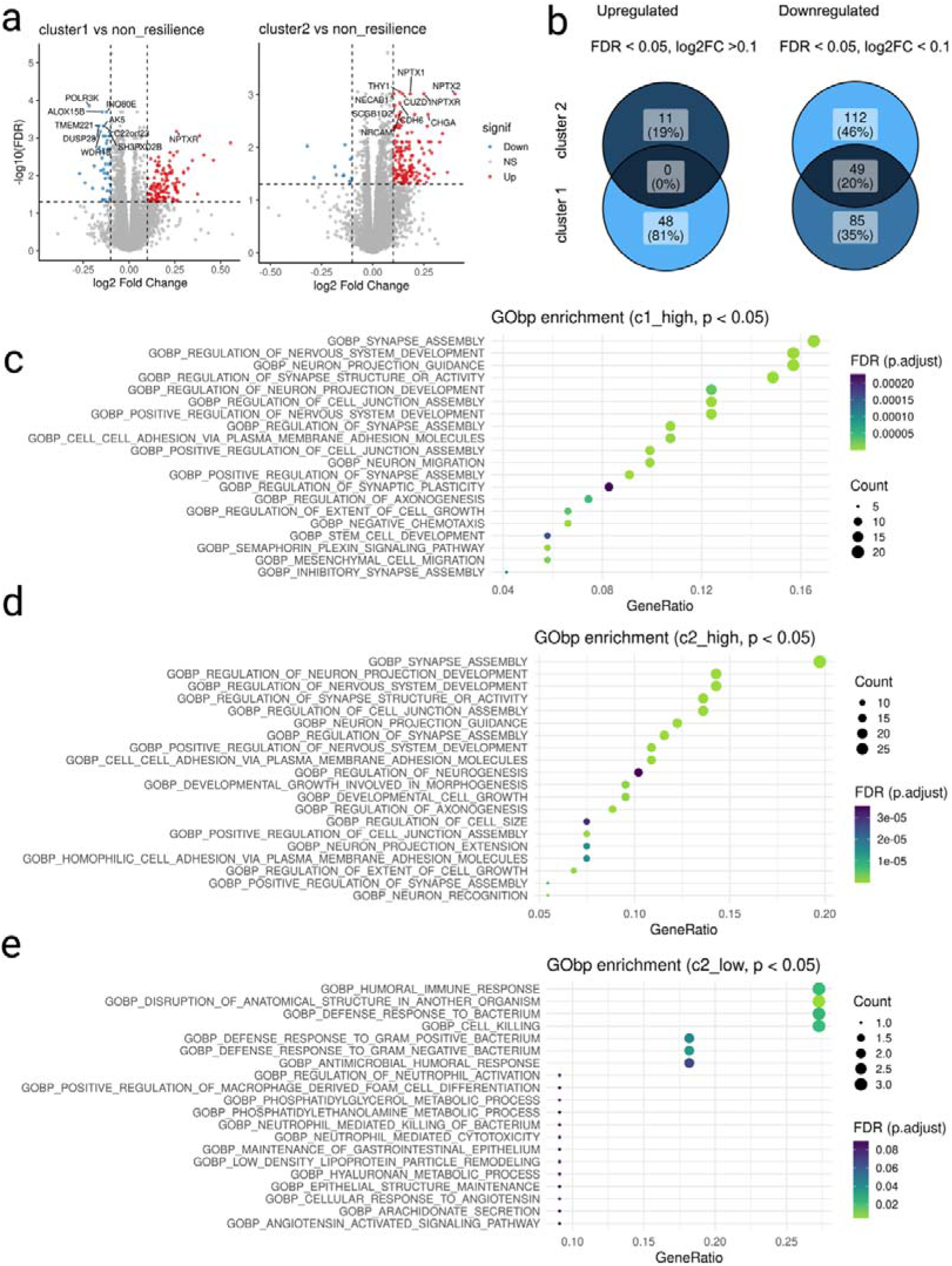
Differential protein expression and pathway enrichment analyses between CR subtypes and the non-CR group. (a) Volcano plots comparing protein expression levels between cluster 1 and non-CR (left) and between cluster 2 and non-CR (right). Proteins with Benjamini–Hochberg (BH)–adjusted p-values < 0.05 and log_₂_ fold change (log_₂_FC) > 0.1 are shown in red, and those with BH-adjusted p-values < 0.05 and log_₂_FC < –0.1 are shown in blue. (b) Venn diagrams showing the overlap of significantly upregulated proteins (BH-adjusted p-value < 0.05, log_₂_FC > 0.1; left) and significantly downregulated proteins (BH-adjusted p-value < 0.05, log_₂_FC < −0.1; right) between the cluster 1 vs. non-CR and cluster 2 vs. non-CR comparisons. (c–e) Top 20 enriched Gene Ontology biological process (GObp) terms based on proteins significantly upregulated or downregulated in each cluster (BH-adjusted p-value < 0.05). (c) Cluster 1–high proteins (log_₂_FC > 0.1); (d) cluster 2–high proteins (log_₂_FC > 0.1); (e) cluster 1–low proteins (log_₂_FC < −0.1).

### Establishment of REGA

Finally, we developed the Resilience Gene Analyzer (REGA), an R Shiny-based web application for visualizing model-derived gene contribution scores for CR prediction (Supplementary Fig. 8). REGA also allows users to visualize the distribution of contribution scores, explore relationships between selected genes and resilience scores, and export summary tables. This tool is accessible at https://igcore.cloud/GerOmics/REsilienceGeneAnalyzer/.

## Discussion

Cognitive resilience (CR) in Alzheimer’s disease (AD) describes the preservation of cognitive function despite substantial AD pathology. Previous studies have identified genetic, transcriptomic, proteomic, and cellular correlates of CR [29], suggesting that resilience may arise through diverse biological processes. However, how these processes are organized into molecularly distinct subtypes and how such heterogeneity relates to clinical and neuroanatomical features remain unclear [1,31]. In this study, we integrated machine learning with multi-cohort transcriptomic, proteomic, and neuroimaging analyses to investigate the molecular architecture of CR. By training CR prediction models in ROSMAP, evaluating their cross-cohort generalizability in MSBB, and projecting ROSMAP/MSBB-derived subtype signatures into ADNI CSF proteomics, we provide evidence that CR includes molecularly heterogeneous subtype-like profiles with partially distinct proteomic characteristics.

A central objective of this study was to use machine learning to extract transcriptomic features informative of CR and to use these features as a basis for downstream molecular subtype analysis. To minimize information leakage, model development and primary performance estimation were conducted using nested cross-validation within ROSMAP, with feature selection, feature-number selection, and hyperparameter tuning confined to the inner cross-validation loop. The resulting models showed modest but consistent predictive performance in ROSMAP and retained moderate performance when applied once to the independent MSBB full cohort. Model discrimination improved in the MSBB top/bottom quartile subset, suggesting that the extracted transcriptomic features may be more informative in individuals with more pronounced resilience phenotypes.

We next used the model-derived positive contributors to investigate molecular heterogeneity among CR individuals. Hierarchical clustering separated CR individuals in the ROSMAP/MSBB RNA-seq data into two molecular subgroups. Because ADNI validation relied on CSF proteomics rather than brain transcriptomics, the cross-platform analysis was limited to 22 overlapping genes/proteins. Nevertheless, this reduced signature retained subtype-associated differences in the ROSMAP/MSBB RNA-seq data and enabled centroid-based projection into ADNI. The resulting ADNI cluster-like groups showed distinct projection scores, and several shared proteins showed directionally consistent differences between the projected groups. These results provide partial but consistent cross-cohort and cross-platform support for the ROSMAP/MSBB-derived subtype structure, while indicating that the ADNI-projected groups should be interpreted as cluster-like molecular profiles rather than exact replications of the original RNA-seq-defined subtypes.

The ADNI imaging and longitudinal cognitive analyses provide an important context for interpreting the molecular subtypes. Although both projected CR clusters showed preserved global brain tissue-volume profiles and slower MMSE decline relative to non-CR individuals, clear structural or longitudinal cognitive differences between the two CR clusters were not observed after multiple-testing correction. Thus, the identified molecular subtypes do not appear to represent clearly separable imaging-defined or trajectory-defined resilience phenotypes. Rather, they may reflect partially distinct molecular profiles associated with a broadly similar resilient phenotype. Consistent with this interpretation, CSF proteomic analyses suggested that cluster 2 was more strongly associated with neuronal and synaptic maintenance, including higher NPTX2 levels, whereas cluster 1 was characterized by lower expression of proteins enriched in immune-related pathways, including inflammatory response and phagocytosis. One possible interpretation is that cluster 2 may reflect a resilience state more closely linked to synaptic maintenance, whereas cluster 1 may reflect a state characterized by relatively reduced neuroimmune activation together with preserved neuronal function. This interpretation remains hypothesis-generating, particularly because CSF protein levels may not directly reflect region-specific brain tissue processes.

This study has several limitations. First, although nested cross-validation was used to reduce information leakage during model development, the sample size for transcriptomic machine learning remained modest relative to the dimensionality of the gene-expression data. Second, CR definitions differed across cohorts because available cognitive and pathological measures were not identical: ROSMAP used MMSE-based resilience scores, MSBB used CDR-based scores, and ADNI used CSF biomarker-based AD pathology with MMSE-based resilience scoring. These differences likely contributed to the attenuated external validation performance in MSBB and should be considered when interpreting cross-cohort generalizability. Third, ROSMAP and MSBB differed in brain-region sampling, with ROSMAP primarily representing dorsolateral prefrontal cortex and MSBB including frontal pole or related frontal-region samples. Although batch correction and PCA were used to reduce major cohort-related variation, these approaches cannot fully separate technical batch effects from biological region-specific expression differences. Thus, some region-dependent CR signals may have been attenuated, and the cross-cohort findings should be interpreted as reproducible signals despite tissue heterogeneity rather than as definitive region-independent signatures. Fourth, ADNI validation used CSF proteomics rather than brain RNA-seq, and only a subset of the ROSMAP/MSBB subtype genes overlapped with measured CSF proteins. Therefore, the ADNI analysis provides cross-platform support rather than direct replication of the brain transcriptomic subtype structure. Fifth, the number of ADNI participants with both CSF proteomics and complete imaging or longitudinal cognitive data was limited, reducing statistical power to detect subtle differences between projected CR subtypes. Finally, the model-derived genes and subtype-associated proteins should be interpreted as candidate correlates of CR, not as proven causal mediators. In conclusion, this study provides evidence that cognitive resilience in AD is molecularly heterogeneous. Machine learning models trained on ROSMAP transcriptomic data identified CR-related gene signatures with moderate cross-cohort generalizability in MSBB. Clustering of model-derived genes revealed two molecular CR subgroups, and a reduced cross-platform signature provided partial support for these subtypes in ADNI CSF proteomics. Both projected CR subtypes showed preserved global brain tissue measures and slower MMSE decline relative to non-CR participants, while clear imaging or longitudinal cognitive differences between the two CR clusters were not observed. These findings suggest that distinct molecular profiles may converge on a shared resilient phenotype in AD and provide a framework for future studies aimed at dissecting the biological mechanisms underlying cognitive resilience.

## Methods

### Data Collection and Preprocessing

#### ROSMAP and MSBB cohort

To construct machine learning models, we used RNA-seq data from the ROSMAP and MSBB cohorts, both of which include comprehensive cognitive assessments as well as pathological evaluations related to amyloid-β and tau. Conditional quantile-normalized expression data were obtained for both cohorts (https://doi.org/10.7303/syn2580853). Details of sample collection and data processing have been described previously [37,38]. For the ROSMAP cohort, samples were derived from the dorsolateral prefrontal cortex or frontal cortex, and sequencing batches 8 and 9 were excluded from the analysis. The MSBB cohort included samples from the frontal pole. Based on the definition of cognitive resilience (CR) as preserved cognitive function in the presence of AD pathology, we included ROSMAP samples with ceradsc scores of 1 or 2 and Braak stage ≥4. In ROSMAP, ceradsc is coded as 1 = Definite AD, 2 = Probable AD, 3 = Possible AD, and 4 = No AD, which is the reverse order of the conventional CERAD C-score coding. For the MSBB cohort, which uses conventional CERAD coding, we included samples with CERAD scores ≥2 and Braak stage ≥4. Ultimately, we analyzed 155 samples from the MSBB cohort and 433 samples from the ROSMAP cohort, including both dementia patients and cognitively unimpaired individuals. Batch effects were corrected using ComBat [61] based on the batch information in each dataset. A total of 18,080 genes that were commonly expressed in both cohorts were included in the analysis.

#### ADNI

For cross-platform evaluation, we used data from the Alzheimer’s Disease Neuroimaging Initiative (ADNI) cohort (adni.loni.usc.edu). The ADNI was launched in 2003 as a public-private partnership, led by Principal Investigator Michael W. Weiner, MD. The original goal of ADNI was to test whether serial magnetic resonance imaging (MRI), positron emission tomography (PET), other biological markers, and clinical and neuropsychological assessment can be combined to measure the progression of mild cognitive impairment (MCI) and early Alzheimer’s disease (AD). The current goals include validating biomarkers for clinical trials, improving the generalizability of ADNI data by increasing diversity in the participant cohort, and to provide data concerning the diagnosis and progression of Alzheimer’s disease to the scientific community. For up-to-date information, see adni.loni.usc.edu. Based on previous studies, AD pathology positivity was defined as a CSF Elecsys immunoassay [62] tTau/Aβ42 ratio > 0.33, pTau/Aβ42 ratio > 0.028, and Aβ42 ≤ 880 pg/mL. CSF proteomics data were obtained using the SOMAscan platform, a high-throughput proteomics technology that utilizes SOMAmer (Slow Off-rate Modified Aptamer) reagents. SOMAmers are synthetic DNA-based aptamers with high affinity and specificity for their protein targets, enabling simultaneous quantification of thousands of proteins in a single assay. We used previously reported SOMAscan data that were normalized and quality-controlled according to established protocols [63]. Only baseline data were used for the analyses.

#### Quantification of CR

A linear regression analysis was performed to calculate CR scores. The score for each individual was defined as the difference between observed and predicted cognitive functions. To predict cognitive function, clinical dementia rating (CDR) was used as the target variable for the MSBB cohort and the mini-mental state examination (MMSE) was used for the ROSMAP and ADNI cohort. The explanatory variables included the CERAD score and Braak stage for ROSMAP and MSBB, and Aβ42, Tau, pTau for ADNI, respectively. As a sensitivity analysis, we recalculated resilience scores after including available demographic covariates. In ROSMAP, age at death, sex, and years of education were included; in MSBB, age at death and sex were included because years of education were not available. These covariate-adjusted scores were compared with the primary pathology-based resilience scores to assess robustness.

For the primary binary CR phenotype, resilience scores were first estimated in the eligible samples of each cohort using the regression models described above. Within the AD pathology–positive analysis set, participants were classified by the sign of the resilience score: scores ≥ 0 were defined as high-CR/CR and scores < 0 as low-CR/non-CR. For MSBB, where higher CDR indicates worse cognitive status, the score was calculated as predicted CDR minus observed CDR so that positive values consistently represented better-than-predicted cognitive status. The MSBB top/bottom quartile analysis was used only as a secondary sensitivity analysis and did not define the primary CR/non-CR groups. Machine Learning Models We evaluated four supervised classifiers: ElasticNet-regularized logistic regression, support vector machine (SVM) [41], Random Forest [42], and XGBoost [43]. ElasticNet [40] and SVM models were implemented using scikit-learn pipelines that included standardization of input features. Random Forest was implemented using RandomForestClassifier, and XGBoost was implemented using XGBClassifier. Hyperparameters were tuned within the inner cross-validation loop of the nested cross-validation framework. The hyperparameter search space and selected configurations are summarized in Supplementary Table S2.

#### Nested Cross-Validation, Feature Selection, and Hyperparameter Tuning

Model development and primary performance estimation were performed in the ROSMAP cohort using nested cross-validation to minimize information leakage. ROSMAP samples were split into five outer folds using stratified cross-validation. In each outer iteration, four folds were used as the outer training set, and the remaining fold was held out as the outer test set. The outer test set was kept completely independent and was not used for feature selection, hyperparameter tuning, feature-number selection, or model selection.

Within each outer training set, model selection was performed using inner 5-fold cross-validation. In each inner fold, feature selection was conducted using only the inner training data. First, ElasticNet-based feature preselection was performed using regularized logistic regression, and genes with non-zero coefficients were retained as candidate features. When the number of selected genes exceeded 500, genes were ranked by the absolute value of their ElasticNet coefficients, and the top 500 genes were retained. Candidate features were then ranked using permutation importance calculated within the corresponding inner training data. Thus, both feature preselection and feature ranking were performed independently within each inner training fold, without using the corresponding inner validation fold or the outer test fold.

For each model, candidate features retained after ElasticNet-based preselection were ranked using permutation importance calculated within the corresponding inner training data. We then evaluated candidate hyperparameter settings together with feature-number settings corresponding to the top 5, 10, 20, 50, 100, 200, or all ranked candidate features. Because the number of ElasticNet-selected candidate features varied across folds, the “all” setting corresponded to all ranked candidate features available within each fold. Candidate configurations were assessed on the corresponding inner validation fold using AUROC. The hyperparameter and feature-number setting yielding the highest mean inner-validation AUROC across the inner validation results was selected as the best configuration for that outer fold.

After model selection, the selected configuration was refitted using the full outer training set. Feature preselection, feature ranking, feature-number selection, and model fitting were repeated using only the full outer training data, and the resulting model was evaluated once on the held-out outer test set. Repeating this procedure across all five outer folds generated out-of-fold predictions for all ROSMAP samples, which were used as the primary estimate of model performance. The hyperparameter search space and selected configurations are summarized in Supplementary Table S2.

Primary model performance was evaluated using the ROSMAP out-of-fold predictions. We calculated AUROC, AUPRC, accuracy, recall, precision, F1 score, and Brier score for each model. Threshold-dependent metrics, including accuracy, recall, precision, and F1 score, were calculated using a probability threshold of 0.5.

For external validation, final models were trained using the full ROSMAP cohort after model configuration had been fixed within ROSMAP. The trained models were then applied once to the independent MSBB full cohort. The MSBB cohort was not used for feature selection, hyperparameter tuning, feature-number selection, or model selection, and was therefore treated as an untouched external validation dataset.

As a secondary sensitivity analysis, we also evaluated the same fixed ROSMAP-trained models in the MSBB top/bottom quartile subset defined by CR score. This analysis was performed only after final model fixation and was not used for any model-selection procedure.

For all performance metrics, 95% confidence intervals were estimated using nonparametric bootstrap resampling with 1,000 iterations. In each iteration, samples were drawn with replacement from the evaluation set, and performance metrics were recalculated using the corresponding true labels and predicted probabilities. Bootstrap samples containing only one class were excluded from the bootstrap summary. The 2.5th and 97.5th percentiles of the bootstrap distribution were used as the lower and upper bounds of the 95% confidence interval.

### Feature Attribution for Biological Interpretation

For ElasticNet, feature importance was determined based on the model coefficients. For SVM, Random Forest, and XGBoost models, SHAP (SHapley Additive exPlanations) values were calculated to quantify feature importance. SHAP values represent the extent to which each feature (gene) contributes to the model output. The SHAP approach is grounded in cooperative game theory and fairly distributes the “payout” among features by considering all possible combinations [39]. We used the Python SHAP library for these analyses: TreeExplainer was applied for Random Forest and XGBoost, and KernelExplainer was used for SVM.

### Hierarchical Clustering

Hierarchical clustering was performed using the pheatmap function in R. For exploratory CR subtype discovery in ROSMAP/MSBB, CR samples were clustered using z-scored expression values of 144 model-derived genes with positive contributions to CR prediction. Gene expression values were scaled by gene, and hierarchical clustering was performed using Euclidean distance and Ward.D2 linkage. Given the limited number of CR samples relative to the number of clustering features, we focused on the primary bifurcation of the hierarchical dendrogram to avoid defining multiple small, potentially unstable clusters. Two major sample clusters were therefore extracted for downstream exploratory analyses and provisionally annotated as CR cluster 1 and CR cluster 2. Because formal cluster-number optimization and bootstrap-based stability assessment were not performed, and because clustering used supervised, model-derived features rather than unsupervised genome-wide features, this two-cluster structure was interpreted as an exploratory candidate subtype structure.

### Imaging Data Acquisition from ADNI cohort

#### Tissue Volume Quantification by WMH

White matter hyperintensity (WMH) volumes were quantified as described previously[64–67]. Briefly, WMH segmentation was performed using a fully automated Bayesian framework based on high-resolution T1-weighted and FLAIR MRI scans. After coregistration and inhomogeneity correction of T1 and FLAIR images, each scan was non-linearly aligned to a common template space. WMH segmentation utilized prior probability maps derived from manually labeled cases, combined with image intensity histogram fitting and thresholding. Probabilistic likelihood values of WMH were calculated for each voxel and thresholded at 3.5 standard deviations above the mean to generate binary WMH masks. The segmented masks were then back-transformed into native space to compute total WMH volume, which was reported in cubic centimeters (cm³). For the quantification of gray matter, white matter, and cerebrospinal fluid (CSF) volumes, tissue segmentation was performed on the native-space T1-weighted images using an Expectation-Maximization (EM) algorithm. This method iteratively refined tissue classifications based on Gaussian models of image intensity and spatial priors derived from template-based masks. To further improve segmentation accuracy, a Markov Random Field (MRF) model was incorporated to enforce spatial smoothness. Final tissue labels were assigned after several iterations, and voxels labeled as WMH were substituted for normal-appearing white matter to generate a four-tissue segmentation (gray matter, white matter, CSF, and WMH). All regional volumes were calculated in native space and reported in cubic centimeters (cm³). The UCD table included CEREBRUM_TCV, defined as total cerebrum cranial volume for the supratentorial compartment. In the primary UCD imaging analyses, CEREBRUM_TCV was used as an intracranial-size proxy for volumetric adjustment. CEREBRUM_TCV itself was not adjusted for CEREBRUM_TCV when analyzed descriptively as an imaging measure.

### Cortical and Subcortical Thickness and Volume Quantification with FreeSurfer

Cortical reconstruction and volumetric segmentation were performed using the FreeSurfer image analysis suite (version 7.1.1; http://surfer.nmr.mgh.harvard.edu/fswiki/FreeSurferMethodsCitation), as previously described[54–60]. Briefly, T1-weighted MRI scans from the ADNI cohort were preprocessed at the Mayo Clinic, including gradient warping, intensity scaling, B1 correction, and N3 inhomogeneity correction. After visual inspection for image quality, the standard cross-sectional FreeSurfer pipeline was applied, incorporating motion correction, skull stripping, intensity normalization, automated Talairach transformation, and segmentation of subcortical white matter and deep gray matter structures. The cortical surface was reconstructed and parcellated into gyral-based regions of interest using the Desikan-Killiany atlas. Only scans rated as “Pass” by visual QC were included in the analysis. FreeSurfer-derived regional volume metrics were obtained from the ADNI UCSF FreeSurfer summary tables and analyzed as supportive supplementary imaging measures. The available regional volume measures included cortical and subcortical volumetric summaries derived from the standard FreeSurfer processing pipeline. Because these analyses were based on released summary tables rather than reprocessing of raw MRI scans, and because a consistent eTIV-equivalent covariate was not available for the final supportive analysis, FreeSurfer results were interpreted as supplementary rather than primary imaging evidence. All volumetric measures were calculated in native space and reported in cubic millimeters (mm³). Because an eTIV-equivalent covariate was not available in the UCSF FreeSurfer table used for this analysis, FreeSurfer regional analyses were treated as supportive supplementary analyses rather than primary imaging evidence. Statistics

All statistical analyses were performed using R (version 4.4.0) and Python (version 3.10) with relevant packages. Unless otherwise specified for multiple comparisons, a two-sided p-value < 0.05 was considered statistically significant. For analyses involving multiple imaging metrics or multiple prespecified pairwise contrasts, p-values were adjusted using the Benjamini–Hochberg procedure within the corresponding analysis family. The primary UCD imaging analysis applied FDR correction across all UCD metrics and all prespecified pairwise contrasts. FreeSurfer regional analyses were reported as supportive supplementary analyses.

### Comparison of Clinical and Imaging Data among Groups

Group differences in demographic variables were evaluated using Kruskal–Wallis tests for continuous variables and chi-squared tests for categorical variables. For the primary UCD imaging analysis, each WMH or global tissue-volume metric was modeled using covariate-adjusted linear regression with projected group as the main predictor. Models were adjusted for age at MRI, sex, years of education, APOE ε4 carrier status, ADNI phase, scanner manufacturer, and magnetic field strength when estimable. For volumetric measures, CEREBRUM_TCV was additionally included as an intracranial-size proxy, except when CEREBRUM_TCV itself was the outcome. TOTAL_WMH was analyzed as log10(TOTAL_WMH + 1) because of its skewed distribution.

Prespecified pairwise contrasts were evaluated for cluster 1 versus non-CR, cluster 2 versus non-CR, and cluster 1 versus cluster 2. P-values were adjusted across all UCD imaging metrics and pairwise contrasts using the Benjamini–Hochberg false discovery rate procedure. Effect sizes were reported as adjusted mean differences and standardized adjusted differences with 95% confidence intervals. Kruskal–Wallis tests followed by Dunn’s post hoc tests were performed as supplementary distribution-free sensitivity analyses. UCSF FreeSurfer regional metrics were summarized as supportive supplementary analyses because an eTIV-equivalent covariate was not available in the released UCSF table used in this study.

### Differential Gene Expression Analysis

To compare gene expression levels between CR cluster 1 and CR cluster 2 in the ROSMAP/MSBB RNA-seq data, we performed two-sided Welch’s t-tests for each subtype-associated gene. For the main clustering analysis, 144 model-derived genes with positive contributions to CR prediction were used. For cross-platform comparison with ADNI CSF proteomics, we additionally summarized the subset of 22 genes that overlapped with proteins measured in the ADNI SOMAscan dataset. Multiple testing correction was performed using the Benjamini–Hochberg procedure.

Gene/protein-wise Association Analysis with Clinical and Pathological Variables Associations between gene or protein expression levels and clinical, demographic, or pathological variables were assessed using gene-wise or protein-wise linear models. For continuous or ordinal variables, each expression value was modeled as a function of the clinical variable of interest with relevant covariates. For categorical variables, full and reduced linear models were compared using nested ANOVA.

### Projection of ROSMAP/MSBB CR Subtype Signatures to ADNI CSF Proteomics

To evaluate whether the CR subtype structure identified in ROSMAP/MSBB RNA-seq data was reflected in ADNI CSF proteomics, we performed centroid-based projection using the 22 genes/proteins shared between the ROSMAP/MSBB subtype signature and the ADNI SOMAscan dataset. First, expression values of the 22 shared genes were z-scored in the ROSMAP/MSBB RNA-seq data, and cluster-specific centroids were calculated as the mean z-scored expression profile for CR cluster 1 and CR cluster 2. ADNI CSF protein abundance values for the corresponding 22 proteins were also z-scored. For each ADNI individual, similarity scores were calculated between the ADNI 22-protein profile and each ROSMAP/MSBB cluster centroid using Pearson correlation. Each ADNI individual was assigned to the cluster-like group with the higher centroid similarity score. As sensitivity analyses, we also calculated projection scores using negative Euclidean distance and marker-delta–weighted scoring. For downstream ADNI analyses, participants with AD pathology and high CR were assigned to cluster 1 or cluster 2 according to the higher centroid-correlation score. Participants with AD pathology and low CR were assigned to the non-CR group. Thus, imaging and longitudinal cognitive analyses compared three projected groups: cluster 1, cluster 2, and non-CR.

### Longitudinal MMSE and Time-to-decline Analyses

Longitudinal cognitive trajectories were evaluated using repeated MMSE measurements. The primary longitudinal analysis used linear mixed-effects models with MMSE score as the outcome and projected group, time in years, and the group-by-time interaction as fixed effects. Models were adjusted for age, sex, years of education, and APOE ε4 carrier status. Participant-level random intercepts were included; random slopes for time were evaluated when estimable. Estimated annual MMSE slopes were obtained for each projected group, and pairwise slope contrasts were evaluated for non-CR versus cluster 1, non-CR versus cluster 2, and cluster 1 versus cluster 2. P-values for pairwise slope contrasts were adjusted using the Benjamini–Hochberg procedure.

As supplementary time-to-decline analyses, Cox proportional hazards models were used to evaluate time to incident MMSE decline, defined as the first observed visit with MMSE < 24. Participants who already had MMSE < 24 at the first available MMSE visit were excluded from the primary incident-event risk set. Cox models were adjusted for age, sex, years of education, and APOE ε4 carrier status. Baseline MMSE, CDR, and clinical diagnosis were not included as covariates because these variables are closely related to the CR phenotype and may constitute overadjustment. Pairwise hazard ratios were estimated for cluster 1 versus non-CR, cluster 2 versus non-CR, and cluster 1 versus cluster 2. Kaplan–Meier curves, when shown, were used only for supplementary visualization.

### Differential Protein Expression Analysis

Protein expression levels between groups were compared using Welch’s two-sample t-test, which does not assume equal variances. Missing values were imputed with the minimum observed value for each protein (a left-censored imputation that may bias apparent group differences toward lower-abundance directions) across all samples. P-values were adjusted for multiple comparisons using the Benjamini–Hochberg method, and proteins with BH-adjusted p-values < 0.05 were deemed significant. Log_₂_ fold changes (log_₂_FC) were calculated as the difference in log_₂_-transformed mean expression values between groups.

### Establishment of REGA

We developed the Resilience Gene Analyzer (REGA), an R Shiny-based web application for visualizing model-derived gene contribution scores for CR prediction. For ElasticNet, signed model coefficients were used as gene-level contribution scores. For SVM, Random Forest, and XGBoost, signed mean SHAP values were used for visualization, while mean absolute SHAP values were retained as unsigned measures of contribution strength. REGA allows users to select a model and one or more genes, visualize the distribution of gene-level contribution scores, examine positive or negative contributions to CR prediction, explore relationships with resilience scores, and export summary tables.

## Declarations

### Ethics approval and consent to participate

Not applicable.

### Consent for publication

Not applicable.

### Availability of data and materials

All the data used in this study are publicly accessible. The ROSMAP and MSBB RNA-seq datasets analyzed in this study are available through the AD Knowledge Portal under the applicable data-use agreements. ADNI clinical, CSF biomarker, proteomic, and MRI-derived data are available from the ADNI database subject to ADNI data-use approval. Processed analysis outputs necessary to reproduce the main figures and tables are provided in the Supplementary Information or are available from the corresponding author upon reasonable request, where permitted by the original data-use agreements.

Codes used for the current analysis are available at: https://github.com/matsui-lab/Resilience-prediction-RNA-seq.

## Authors’ contributions

A.K. conceived the study, performed data collection, analysis, and visualization, and wrote the original draft of the manuscript. Y.M. supervised the project, advised on analysis strategies, provided resources, secured funding, and contributed to reviewing and editing the manuscript. All authors read and approved the final manuscript.

## Acknowledgements

The data available in the AD Knowledge Portal would not be possible without the participation of research volunteers and the contribution of data by collaborating researchers. The results published in this study are, in whole or in part, based on data obtained from the AD Knowledge Portal (https://adknowledgeportal.org). Data generation was supported by the following NIH grants: P30AG10161, P30AG72975, R01AG15819, R01AG17917, R01AG036836, U01AG46152, U01AG61356, U01AG046139, P50 AG016574, R01 AG032990, U01AG046139, R01AG018023, U01AG006576, U01AG006786, R01AG025711, R01AG017216, R01AG003949, R01NS080820, U24NS072026, P30AG19610, U01AG046170, RF1AG057440, and U24AG061340. We also acknowledge support from the Cure PSP, Mayo and Michael J Fox foundations, the Arizona Department of Health Services, and the Arizona Biomedical Research Commission. We extend our gratitude to the participants of the Religious Order Study and Memory and Aging projects for their generous donations. Additionally, we thank the Sun Health Research Institute Brain and Body Donation Program, Mayo Clinic Brain Bank, and Mount Sinai/JJ Peters VA Medical Center NIH Brain and Tissue Repository. Data and analysis contributors include Nilüfer Ertekin-Taner, Steven Younkin (Mayo Clinic, Jacksonville, FL), Todd Golde (University of Florida), Nathan Price (Institute for Systems Biology), David Bennett, Christopher Gaiteri (Rush University), Philip De Jager (Columbia University), Bin Zhang, Eric Schadt, Michelle Ehrlich, Vahram Haroutunian, Sam Gandy (Icahn School of Medicine at Mount Sinai), Koichi Iijima (National Center for Geriatrics and Gerontology, Japan), Scott Noggle (New York Stem Cell Foundation), and Lara Mangravite (Sage Bionetworks). The grantee organization is the Northern California Institute for Research and Education. In the past, ADNI has also received funding from the National Institute of Biomedical Imaging and Bioengineering, the Canadian Institutes of Health Research, and private sector contributions through the Foundation for the National Institutes of Health (FNIH) including generous contributions from the following: AbbVie, Alzheimer’s Association; Alzheimer’s Drug Discovery Foundation; Araclon Biotech; BioClinica, Inc.; Biogen; Bristol-Myers Squibb Company; CereSpir, Inc.; Cogstate; Eisai Inc.; Elan Pharmaceuticals, Inc.; Eli Lilly and Company; EuroImmun; F. Hoffmann-La Roche Ltd and its affiliated company Genentech, Inc.; Fujirebio; GE Healthcare; IXICO Ltd.; Janssen Alzheimer Immunotherapy Research & Development, LLC.; Johnson & Johnson Pharmaceutical Research & Development LLC.; Lumosity; Lundbeck; Merck & Co., Inc.; Meso Scale Diagnostics, LLC.; NeuroRx Research; Neurotrack Technologies; Novartis Pharmaceuticals Corporation; Pfizer Inc.; Piramal Imaging; Servier; Takeda Pharmaceutical Company; and Transition Therapeutics. Finally, all figures in this study were created using BioRender.com.

## List of Abbreviations

Aβ: amyloid-beta
AD: Alzheimer’s disease
ADNI: Alzheimer’s Disease Neuroimaging Initiative
AUPRC: area under the precision–recall curve
AUROC: area under the receiver operating characteristic curve
CDR: Clinical Dementia Rating
CERAD: Consortium to Establish a Registry for Alzheimer’s Disease
CR: cognitive resilience
CSF: cerebrospinal fluid
FDR: false discovery rate
MMSE: Mini-Mental State Examination
MRI: magnetic resonance imaging
MSBB: Mount Sinai Brain Bank
ROSMAP: Religious Orders Study and Memory and Aging Project
SHAP: Shapley additive explanations
WMH: white matter hyperintensity.

## Additional files

Supplementary Figures – word file

Supplementary Fig. 1-8

Supplementary Table S1.

Supplementary Table S2.

Supplementary Table S3.

Supplementary Table S4.

Supplementary Table S5.

Supplementary Table S6.

Supplementary Table S7.

Supplementary Table S8.

Supplementary Table S9.

Supplementary Table S10.

## References

1. de Vries LE, Huitinga I, Kessels HW, Swaab DF, Verhaagen J. The concept of resilience to Alzheimer’s Disease: current definitions and cellular and molecular mechanisms. Mol Neurodegener. 2024;19:33.

2. Negro D, Opazo P. Cognitive resilience in Alzheimer’s disease: from large-scale brain networks to synapses. Brain Commun. 2024;6:fcae050.

3. Arenaza-Urquijo EM, Vemuri P. Resistance vs resilience to Alzheimer disease: Clarifying terminology for preclinical studies. Neurology. 2018;90:695–703.

4. Arenaza-Urquijo EM, Boyle R, Casaletto K, Anstey KJ, Vila-Castelar C, Colverson A, et al. Sex and gender differences in cognitive resilience to aging and Alzheimer’s disease. Alzheimers Dement [Internet]. 2024; Available from: https://alz-journals.onlinelibrary.wiley.com/doi/10.1002/alz.13844

5. Aiello Bowles EJ, Crane PK, Walker RL, Chubak J, LaCroix AZ, Anderson ML, et al. Cognitive resilience to Alzheimer’s disease pathology in the human brain. J Alzheimers Dis. 2019;68:1071–83.

6. Ossenkoppele R, Lyoo CH, Jester-Broms J, Sudre CH, Cho H, Ryu YH, et al. Assessment of demographic, genetic, and imaging variables associated with brain resilience and cognitive resilience to pathological tau in patients with Alzheimer disease. JAMA Neurol. 2020;77:632–42.

7. Carrigan M, Bocancea DI, Vogel J, van Loenhoud AC, Tesi N, Barkhof F, et al. A novel approach to resilience and its links with education and Alzheimer’s disease genetics. Alzheimers Dement. 2025;21:e70379.

8. Willroth EC, James BD, Graham EK, Kapasi A, Bennett DA, Mroczek DK. Well-being and cognitive resilience to dementia-related neuropathology. Psychol Sci. 2023;34:283–97.

9. Yao T, Sweeney E, Nagorski J, Shulman JM, Allen GI. Quantifying cognitive resilience in Alzheimer’s Disease: The Alzheimer’s Disease Cognitive Resilience Score. PLoS One. 2020;15:e0241707.

10. Bocancea DI, Svenningsson AL, van Loenhoud AC, Groot C, Barkhof F, Strandberg O, et al. Determinants of cognitive and brain resilience to tau pathology: a longitudinal analysis. Brain. 2023;146:3719–34.

11. Neuner SM, Telpoukhovskaia M, Menon V, O’Connell KMS, Hohman TJ, Kaczorowski CC. Translational approaches to understanding resilience to Alzheimer’s disease. Trends Neurosci. 2022;45:369–83.

12. Mathys H, Boix CA, Akay LA, Xia Z, Davila-Velderrain J, Ng AP, et al. Single-cell multiregion dissection of Alzheimer’s disease. Nature. 2024;1–11.

13. Berson E, Sreenivas A, Phongpreecha T, Perna A, Grandi FC, Xue L, et al. Whole genome deconvolution unveils Alzheimer’s resilient epigenetic signature. Nat Commun. 2023;14:4947.

14. Tan MS, Cheah P-L, Chin A-V, Looi L-M, Chang S-W. A review on omics-based biomarkers discovery for Alzheimer’s disease from the bioinformatics perspectives: Statistical approach vs machine learning approach. Comput Biol Med. 2021;139:104947.

15. Lopera F, Marino C, Chandrahas AS, O’Hare M, Villalba-Moreno ND, Aguillon D, et al. Resilience to autosomal dominant Alzheimer’s disease in a Reelin-COLBOS heterozygous man. Nat Med. 2023;29:1243–52.

16. Phongpreecha T, Godrich D, Berson E, Espinosa C, Kim Y, Cholerton B, et al. Quantitative estimate of cognitive resilience and its medical and genetic associations. Alzheimers Res Ther. 2023;15:192.

17. Arboleda-Velasquez JF, Lopera F, O’Hare M, Delgado-Tirado S, Marino C, Chmielewska N, et al. Resistance to autosomal dominant Alzheimer’s disease in an APOE3 Christchurch homozygote: a case report. Nat Med. 2019;25:1680–3.

18. Franzmeier N, Ren J, Damm A, Monté-Rubio G, Boada M, Ruiz A, et al. The BDNFVal66Met SNP modulates the association between beta-amyloid and hippocampal disconnection in Alzheimer’s disease. Mol Psychiatry. 2021;26:614–28.

19. Vélez JI, Chandrasekharappa SC, Henao E, Martinez AF, Harper U, Jones M, et al. Pooling/bootstrap-based GWAS (pbGWAS) identifies new loci modifying the age of onset in PSEN1 p.Glu280Ala Alzheimer’s disease. Mol Psychiatry. 2013;18:568–75.

20. Ramanan VK, Lesnick TG, Przybelski SA, Heckman MG, Knopman DS, Graff-Radford J, et al. Coping with brain amyloid: genetic heterogeneity and cognitive resilience to Alzheimer’s pathophysiology. Acta Neuropathol Commun. 2021;9:48.

21. Dumitrescu L, Mahoney ER, Mukherjee S, Lee ML, Bush WS, Engelman CD, et al. Genetic variants and functional pathways associated with resilience to Alzheimer’s disease. Brain. 2020;143:2561–75.

22. Belloy ME, Napolioni V, Han SS, Le Guen Y, Greicius MD, Alzheimer’s Disease Neuroimaging Initiative. Association of klotho-VS heterozygosity with risk of Alzheimer disease in individuals who carry APOE4. JAMA Neurol. 2020;77:849–62.

23. Yu L, Tasaki S, Schneider JA, Arfanakis K, Duong DM, Wingo AP, et al. Cortical proteins associated with cognitive resilience in community-dwelling older persons. JAMA Psychiatry. 2020;77:1172–80.

24. Carlyle BC, Kandigian SE, Kreuzer J, Das S, Trombetta BA, Kuo Y, et al. Synaptic proteins associated with cognitive performance and neuropathology in older humans revealed by multiplexed fractionated proteomics. Neurobiol Aging. 2021;105:99–114.

25. Barker SJ, Raju RM, Milman NEP, Wang J, Davila-Velderrain J, Gunter-Rahman F, et al. MEF2 is a key regulator of cognitive potential and confers resilience to neurodegeneration. Sci Transl Med. 2021;13:eabd7695.

26. Barroeta-Espar I, Weinstock LD, Perez-Nievas BG, Meltzer AC, Siao Tick Chong M, Amaral AC, et al. Distinct cytokine profiles in human brains resilient to Alzheimer’s pathology. Neurobiol Dis. 2019;121:327–37.

27. Huang Z, Merrihew GE, Larson EB, Park J, Plubell D, Fox EJ, et al. Brain proteomic analysis implicates actin filament processes and injury response in resilience to Alzheimer’s disease. Nat Commun. 2023;14:2747.

28. Yu L, Petyuk VA, Gaiteri C, Mostafavi S, Young-Pearse T, Shah RC, et al. Targeted brain proteomics uncover multiple pathways to Alzheimer’s dementia. Ann Neurol. 2018;84:78–88.

29. Oh HS-H, Urey DY, Karlsson L, Zhu Z, Shen Y, Farinas A, et al. A cerebrospinal fluid synaptic protein biomarker for prediction of cognitive resilience versus decline in Alzheimer’s disease. Nat Med. 2025;31:1592–603.

30. Jia B, Xu Y, Zhu X. Cognitive resilience in Alzheimer’s disease: Mechanism and potential clinical intervention. Ageing Res Rev. 2025;106:102711.

31. Chew CS, Lee JY, Ng KY, Koh RY, Chye SM. Resilience mechanisms underlying Alzheimer’s disease. Metab Brain Dis. 2025;40:86.

32. Al Olaimat M, Martinez J, Saeed F, Bozdag S, Alzheimer’s Disease Neuroimaging Initiative. PPAD: A deep learning architecture to predict progression of Alzheimer’s disease. bioRxivorg [Internet]. 2023; Available from: https://www.ncbi.nlm.nih.gov/pmc/articles/PMC9915480/

33. Eteleeb AM, Novotny BC, Tarraga CS, Sohn C, Dhungel E, Brase L, et al. Brain high-throughput multi-omics data reveal molecular heterogeneity in Alzheimer’s disease. PLoS Biol. 2024;22:e3002607.

34. Park C, Ha J, Park S. Prediction of Alzheimer’ s disease based on deep neural network by integrating gene expression and DNA methylation dataset. Expert Syst Appl. 2020;140:112873.

35. Beebe-Wang N, Celik S, Weinberger E, Sturmfels P, De Jager PL, Mostafavi S, et al. Unified AI framework to uncover deep interrelationships between gene expression and Alzheimer’s disease neuropathologies. Nat Commun. 2021;12:5369.

36. Rodriguez S, Hug C, Todorov P, Moret N, Boswell SA, Evans K, et al. Machine learning identifies candidates for drug repurposing in Alzheimer’s disease. Nat Commun. 2021;12:1033.

37. Ahammad I, Lamisa AB, Bhattacharjee A, Jamal TB, Arefin MS, Chowdhury ZM, et al. AITeQ: a machine learning framework for Alzheimer’s prediction using a distinctive five-gene signature. Brief Bioinform [Internet]. 2024;25. Available from: https://pubmed.ncbi.nlm.nih.gov/38877887/

38. Pandey D, Onkara Perumal P. A scoping review on deep learning for next-generation RNA-Seq. data analysis. Funct Integr Genomics. 2023;23:134.

39. Deshpande D, Chhugani K, Chang Y, Karlsberg A, Loeffler C, Zhang J, et al. RNA-seq data science: From raw data to effective interpretation. Front Genet [Internet]. 2023;14. Available from: https://www.frontiersin.org/articles/10.3389/fgene.2023.997383/full

40. De Jager PL, Ma Y, McCabe C, Xu J, Vardarajan BN, Felsky D, et al. A multi-omic atlas of the human frontal cortex for aging and Alzheimer’s disease research. Sci Data. 2018;5:180142.

41. Wang M, Beckmann ND, Roussos P, Wang E, Zhou X, Wang Q, et al. The Mount Sinai cohort of large-scale genomic, transcriptomic and proteomic data in Alzheimer’s disease. Sci Data. 2018;5:180185.

42. Lundberg SM, Lee S-I. A unified approach to interpreting model predictions. Neural Inf Process Syst. 2017;4765–74.

43. Zou H, Hastie T. Regularization and variable selection via the elastic net. J R Stat Soc Series B Stat Methodol. 2005;67:301–20.

44. Boser BE, Guyon IM, Vapnik VN. A training algorithm for optimal margin classifiers. Proceedings of the fifth annual workshop on Computational learning theory [Internet]. New York, NY, USA: ACM; 1992. Available from: 10.1145/130385.130401

45. Breiman L. Random Forests | Machine Learning. Mach Learn. 2001;45:5–32.

46. Chen T, Guestrin C. XGBoost: A Scalable Tree Boosting System. Proceedings of the 22nd ACM SIGKDD International Conference on Knowledge Discovery and Data Mining [Internet]. New York, NY, USA: ACM; 2016. Available from: https://dl.acm.org/doi/10.1145/2939672.2939785

47. Nikhil K, Viccaro K, Shah K. Multifaceted regulation of ALDH1A1 by Cdk5 in Alzheimer’s disease pathogenesis. Mol Neurobiol. 2019;56:1366–90.

48. Tao W, Yu L, Shu S, Liu Y, Zhuang Z, Xu S, et al. MiR-204-3p/Nox4 mediates memory deficits in a mouse model of Alzheimer’s disease. Mol Ther. 2021;29:396–408.

49. Swarnkar S, Avchalumov Y, Espadas I, Grinman E, Liu X-A, Raveendra BL, et al. Molecular motor protein KIF5C mediates structural plasticity and long-term memory by constraining local translation. Cell Rep. 2021;36:109369.

50. Durairajan SSK, Selvarasu K, Singh AK, Patnaik S, Iyaswamy A, Jaiswal Y, et al. Unraveling the interplay of kinesin-1, tau, and microtubules in neurodegeneration associated with Alzheimer’s disease. Front Cell Neurosci. 2024;18:1432002.

51. Zhou J, Yang Y, Liu S, Chen J, Liao H, Liang W, et al. Microglial serine racemase knockout alleviates Alzheimer-like neuropathology and behavioral deficit via lactylation-mediated anti-inflammation. Commun Biol. 2026;9:493.

52. Rong X, Yao X, Fang H, Saji J, Qian Y, Gu J-X, et al. Dual PDE4/10A inhibition restores CREB1 function and enhances neuronal resilience in models of alzheimer’s disease. Alzheimers Res Ther. 2025;17:231.

53. Rehker J, Rodhe J, Nesbitt RR, Boyle EA, Martin BK, Lord J, et al. Caspase-8, association with Alzheimer’s Disease and functional analysis of rare variants. PLoS One. 2017;12:e0185777.

54. Li SY, Johnson R, Smyth LC, Dragunow M. Platelet-derived growth factor signalling in neurovascular function and disease. Int J Biochem Cell Biol. 2022;145:106187.

55. Mao R, Zhao S, Chen J, Wang L, Zheng K, Li B. Identification of novel biomarkers for cognitive function via an integrative analysis. J Alzheimers Dis. 2025;108:931–47.

56. Rahman MS, Harrison E, Biggs H, Seikus C, Elliott P, Breen G, et al. Dynamics of cognitive variability with age and its genetic underpinning in NIHR BioResource Genes and Cognition cohort participants. Nat Med. 2024;30:1739–48.

57. Dale AM, Fischl B, Sereno MI. Cortical surface-based analysis. I. Segmentation and surface reconstruction. Neuroimage. 1999;9:179–94.

58. Fischl B, Sereno MI, Dale AM. Cortical surface-based analysis. II: Inflation, flattening, and a surface-based coordinate system. Neuroimage. 1999;9:195–207.

59. Fischl B, Dale AM. Measuring the thickness of the human cerebral cortex from magnetic resonance images. Proc Natl Acad Sci U S A. 2000;97:11050–5.

60. Fischl B, van der Kouwe A, Destrieux C, Halgren E, Ségonne F, Salat DH, et al. Automatically parcellating the human cerebral cortex. Cereb Cortex. 2004;14:11–22.

61. Desikan RS, Ségonne F, Fischl B, Quinn BT, Dickerson BC, Blacker D, et al. An automated labeling system for subdividing the human cerebral cortex on MRI scans into gyral based regions of interest. Neuroimage. 2006;31:968–80.

62. Fischl B, Salat DH, Busa E, Albert M, Dieterich M, Haselgrove C, et al. Whole brain segmentation: automated labeling of neuroanatomical structures in the human brain. Neuron. 2002;33:341–55.

63. Fischl B, Salat DH, van der Kouwe AJW, Makris N, Ségonne F, Quinn BT, et al. Sequence-independent segmentation of magnetic resonance images. Neuroimage. 2004;23 Suppl 1:S69–84.

64. Johnson WE, Li C, Rabinovic A. Adjusting batch effects in microarray expression data using empirical Bayes methods. Biostatistics. 2007;8:118–27.

65. Hansson O, Seibyl J, Stomrud E, Zetterberg H, Trojanowski JQ, Bittner T, et al. CSF biomarkers of Alzheimer’s disease concord with amyloid-β PET and predict clinical progression: A study of fully automated immunoassays in BioFINDER and ADNI cohorts. Alzheimers Dement. 2018;14:1470–81.

66. Timsina J, Gomez-Fonseca D, Wang L, Do A, Western D, Alvarez I, et al. Comparative analysis of Alzheimer’s disease cerebrospinal fluid biomarkers measurement by multiplex SOMAscan platform and immunoassay-based approach. J Alzheimers Dis. 2022;89:193–207.

67. Schwarz C, Fletcher E, DeCarli C, Carmichael O. Fully-automated white matter hyperintensity detection with anatomical prior knowledge and without FLAIR. Inf Process Med Imaging. 2009;21:239–51.

68. DeCarli C, Fletcher E, Ramey V, Harvey D, Jagust WJ. Anatomical mapping of white matter hyperintensities (WMH): exploring the relationships between periventricular WMH, deep WMH, and total WMH burden: Exploring the relationships between periventricular WMH, deep WMH, and total WMH burden. Stroke. 2005;36:50–5.

69. Yoshita M, Fletcher E, Harvey D, Ortega M, Martinez O, Mungas DM, et al. Extent and distribution of white matter hyperintensities in normal aging, MCI, and AD. Neurology. 2006;67:2192–8.

70. Jack CR Jr, Bernstein MA, Fox NC, Thompson P, Alexander G, Harvey D, et al. The Alzheimer’s Disease Neuroimaging Initiative (ADNI): MRI methods. J Magn Reson Imaging. 2008;27:685–91.

