## Supplementary material for "Candidate Molecular Subtypes of Cognitive Resilience in Alzheimer’s Disease: A Multi-Cohort Machine Learning and Neuroimaging Study": supplementary_information.docx

*Data used in preparation of this article were obtained from the Alzheimer's Disease Neuroimaging Initiative (ADNI) database (adni.loni.usc.edu). As such, the investigators within the ADNI contributed to the design and implementation of ADNI and/or provided data but did not participate in the analysis or writing of this report. A complete listing of ADNI investigators can be found at: http://adni.loni.usc.edu/wp-content/uploads/how_to_apply/ADNI_Acknowledgement_List.pdf

^†^Corresponding author

Yusuke Matsui

### Supplementary Fig. 1 Calculation and sensitivity analysis of cognitive resilience scores.

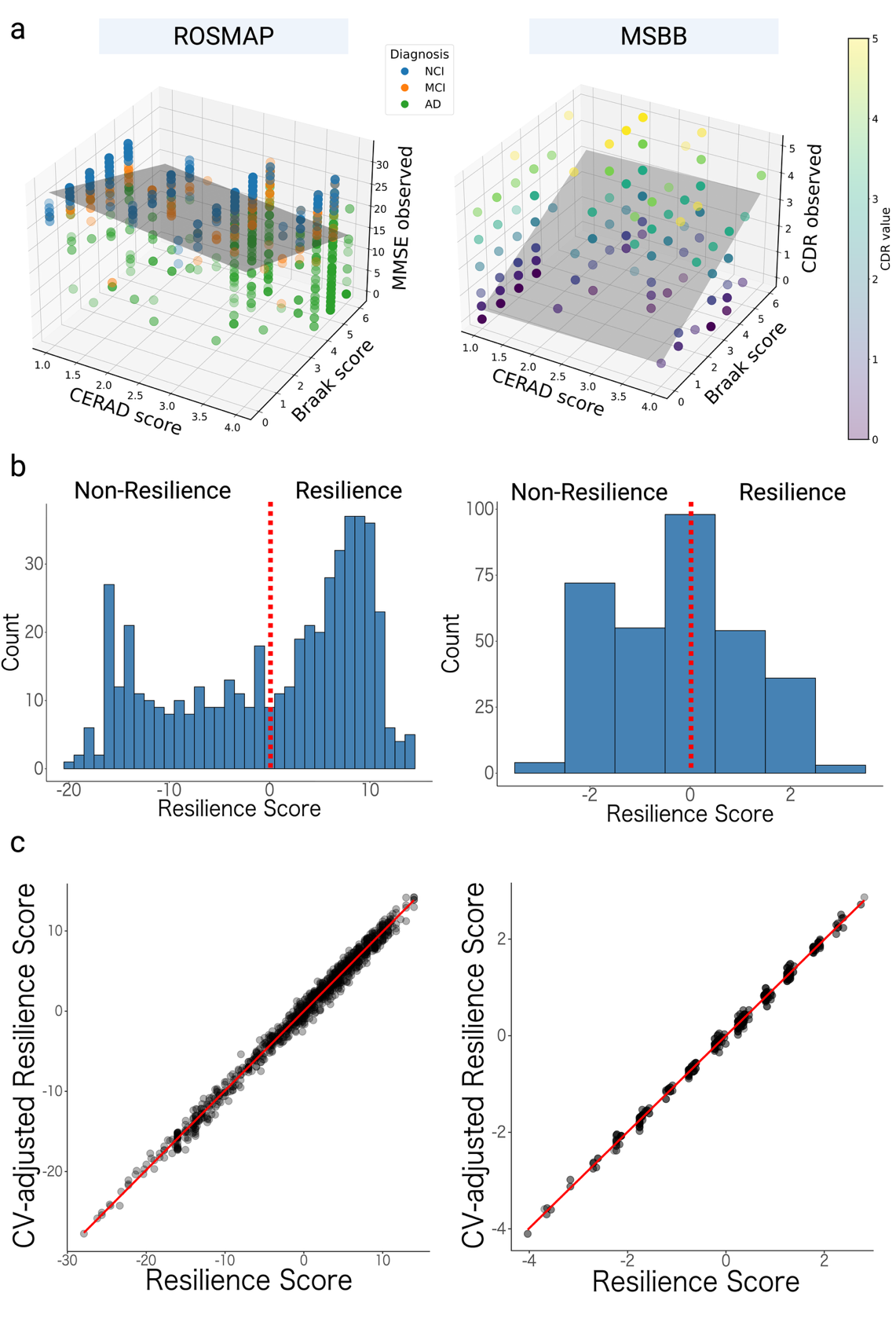

(a) Plots of CERAD score, Braak stage, and cognitive function (MMSE for ROSMAP, CDR for MSBB) with linear regression planes. (b) Histograms of resilience scores. (c) Comparison of resilience scores calculated from pathology-only models and models additionally adjusted for age, sex, and years of education.

### Supplementary Fig. 2 Principal component analysis before batch-effect correction.

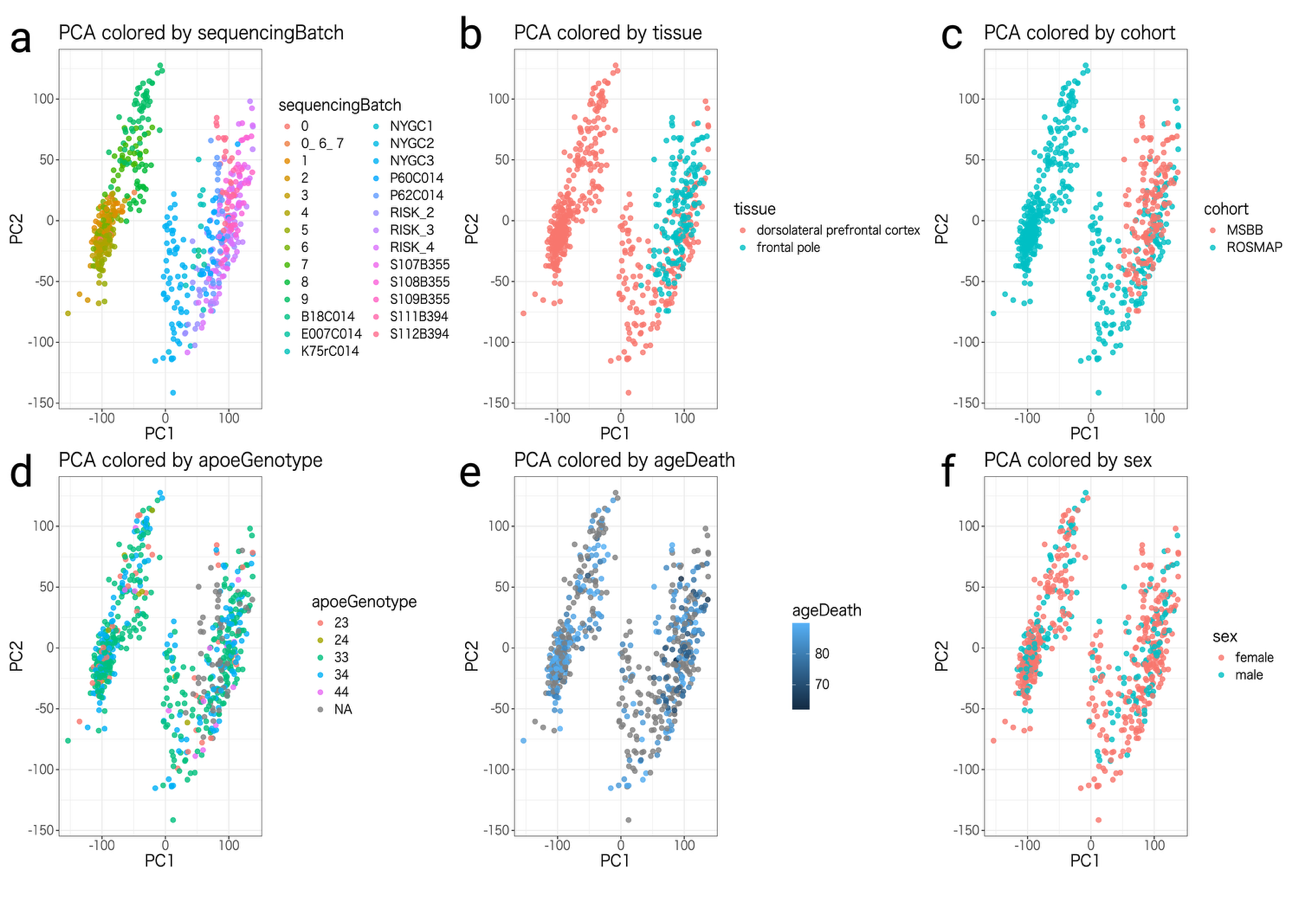

PCA plots of ROSMAP and MSBB RNA-seq data before batch-effect correction, colored by (a) sequencing batch, (b) tissue, (c) cohort, (d) APOE genotype, (e) age, and (f) sex.

### Supplementary Fig. 3 Principal component analysis after batch-effect correction.

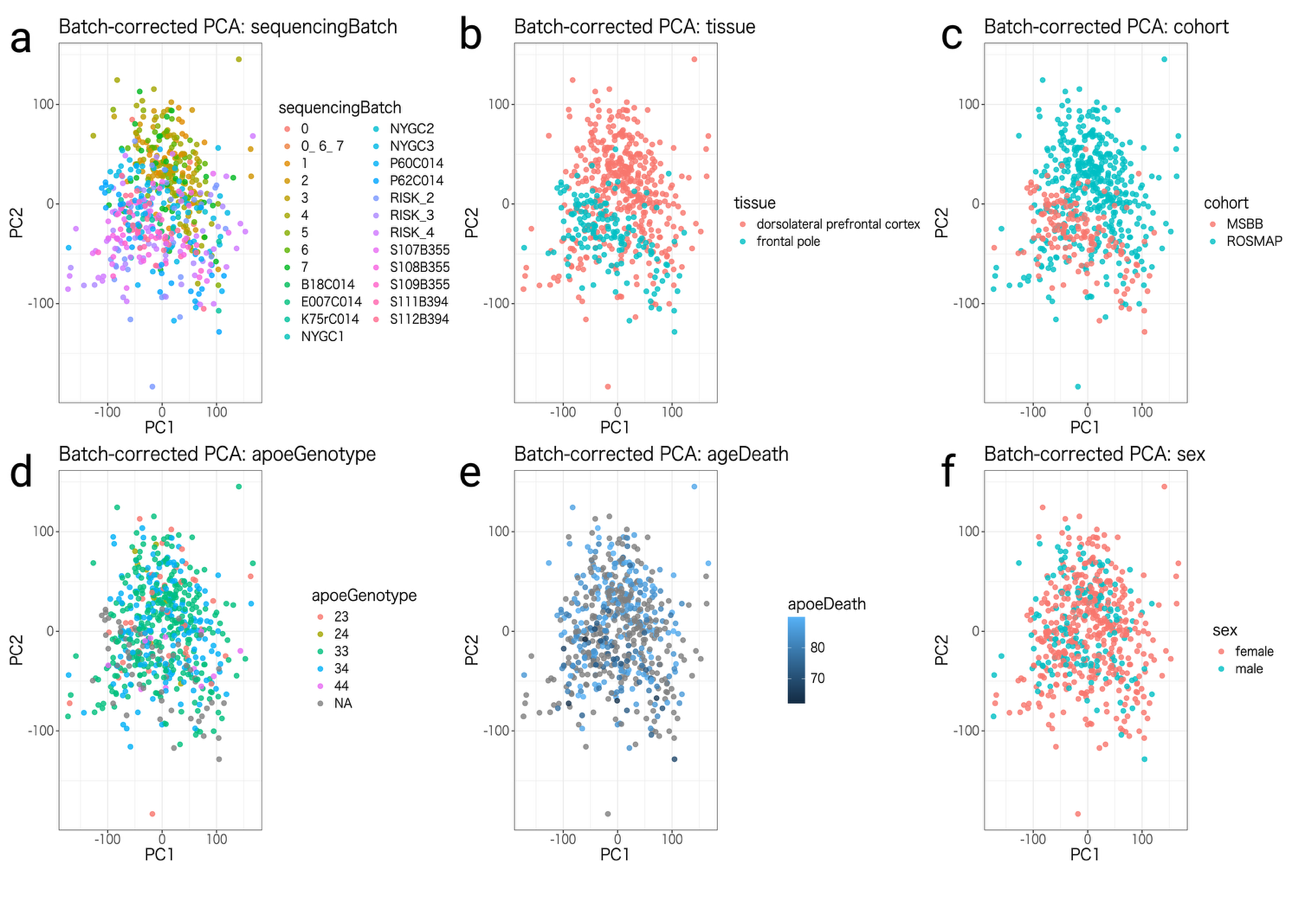

PCA plots of ROSMAP and MSBB RNA-seq data after batch-effect correction, colored by (a) sequencing batch, (b) tissue, (c) cohort, (d) APOE genotype, (e) age, and (f) sex.

### Supplementary Fig. 4 Nested cross-validation and cross-cohort external evaluation workflow.

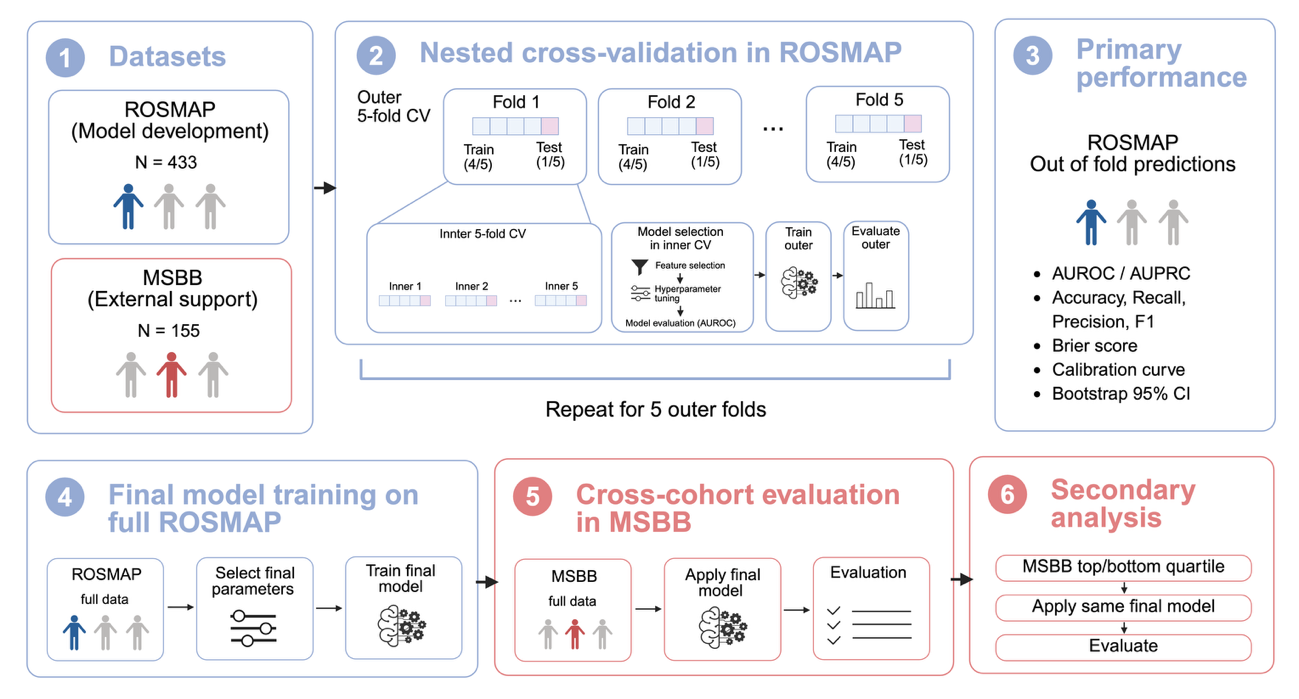

ROSMAP was used for model development and primary performance estimation, whereas MSBB was used only for independent cross-cohort evaluation. In ROSMAP, predictive performance was estimated using nested cross-validation with an outer 5-fold loop and an inner 5-fold loop. Within each outer training set, feature selection, hyperparameter tuning, and feature-number selection were performed only within the inner cross-validation loop. The selected configuration was then refitted using the full outer training set and evaluated once on the held-out outer test fold. Repeating this procedure across all five outer folds generated out-of-fold predictions for all ROSMAP samples, which were used for primary performance evaluation. After this evaluation, final models were trained on the full ROSMAP cohort and applied once to the independent MSBB full cohort. The MSBB top/bottom quartile analysis was performed as a secondary analysis after final model fixation.

### Supplementary Fig. 5 Full performance metrics of CR classification models across ROSMAP and MSBB analyses.

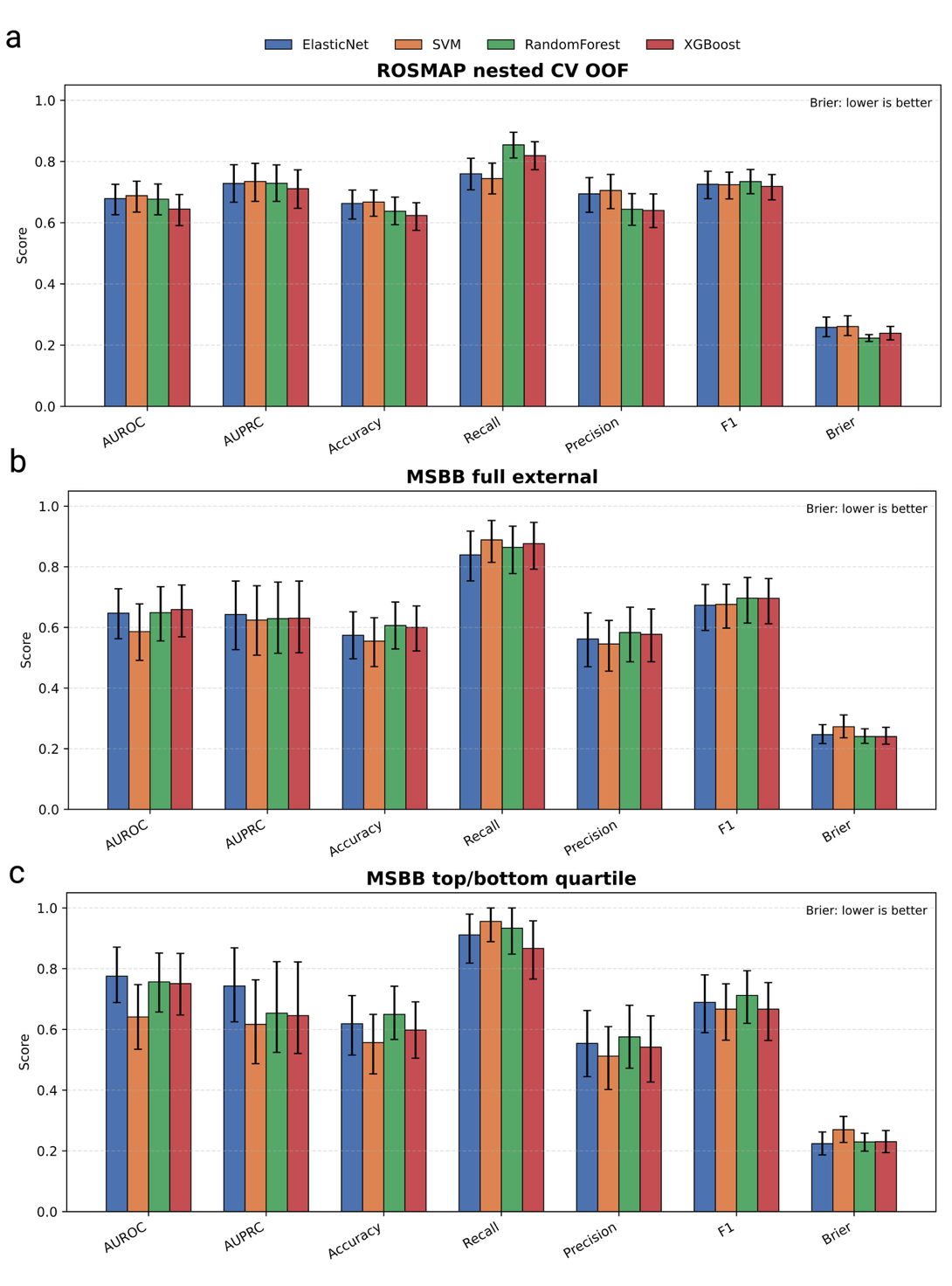

(a) Performance based on out-of-fold predictions from nested cross-validation in ROSMAP. (b) Independent evaluation performance in the full MSBB cohort. (c) Secondary sensitivity analysis in the MSBB top/bottom quartile subset defined by CR score. Bars indicate point estimates, and error bars indicate bootstrap 95% confidence intervals.

### Supplementary Fig. 6. Top model-derived genes contributing positively or negatively to CR prediction.

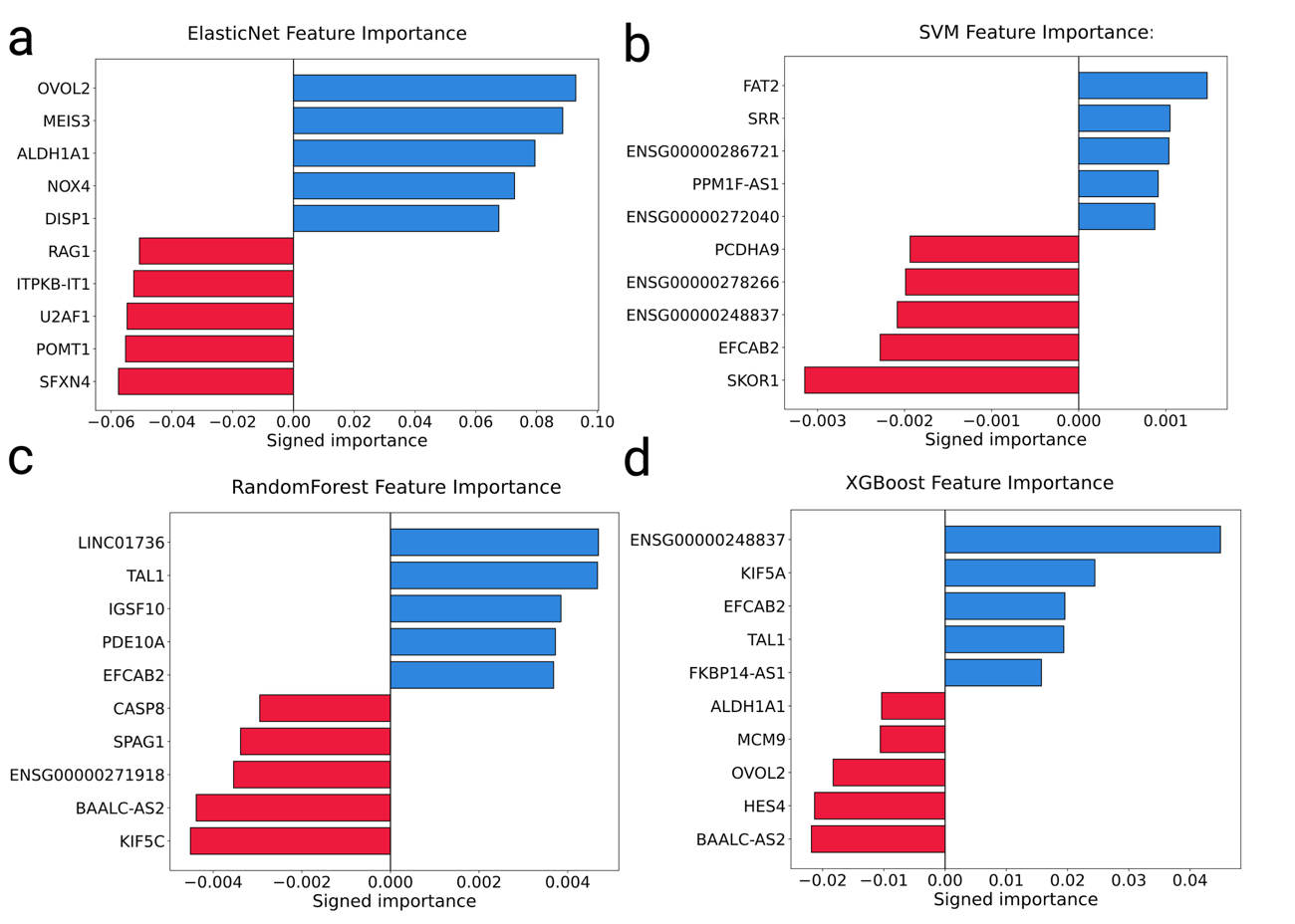

Genes with the strongest positive and negative model-specific contributions to CR prediction are shown for (a) ElasticNet, (b) SVM, (c) Random Forest, and (d) XGBoost models. Positive values indicate contributions toward CR prediction, whereas negative values indicate contributions toward non-CR prediction.

### Supplementary Fig. 7. Clinical Characteristics among CR clusters and non-CR individuals in ADNI cohort.

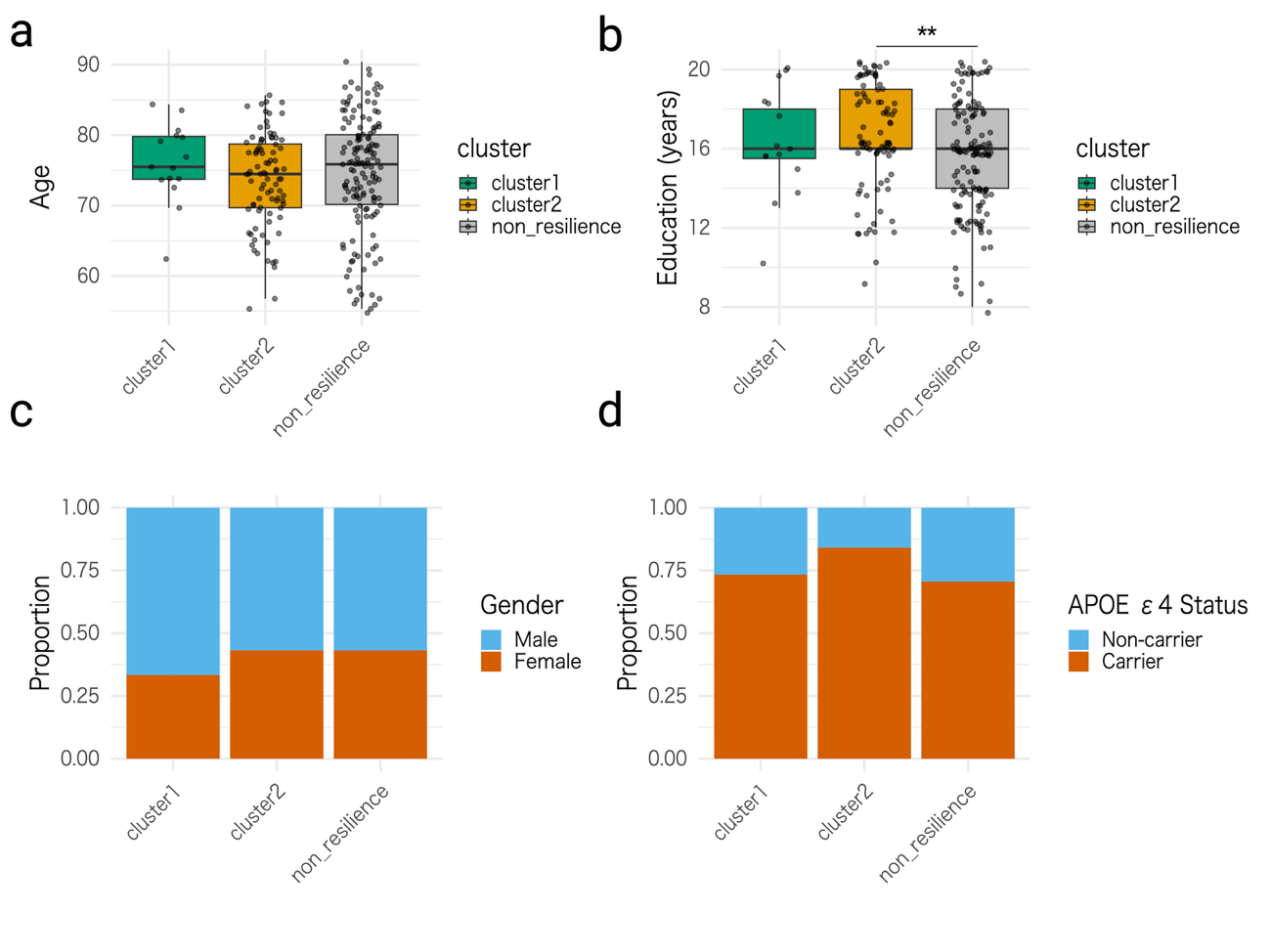

Comparison of (a) age at CSF sampling, (b) years of education, (c) sex distribution, and (d) APOE genotype among Cluster 1, Cluster 2, and non-CR groups in the ADNI cohort.

Statistical significance was assessed using the Kruskal–Wallis test with post hoc Dunn’s test for continuous variables and the chi-squared test for sex and genotype distribution. (**: p < 0.01).

### Supplementary Fig. 8. Visualization of gene-level contributions to CR prediction using REGA.

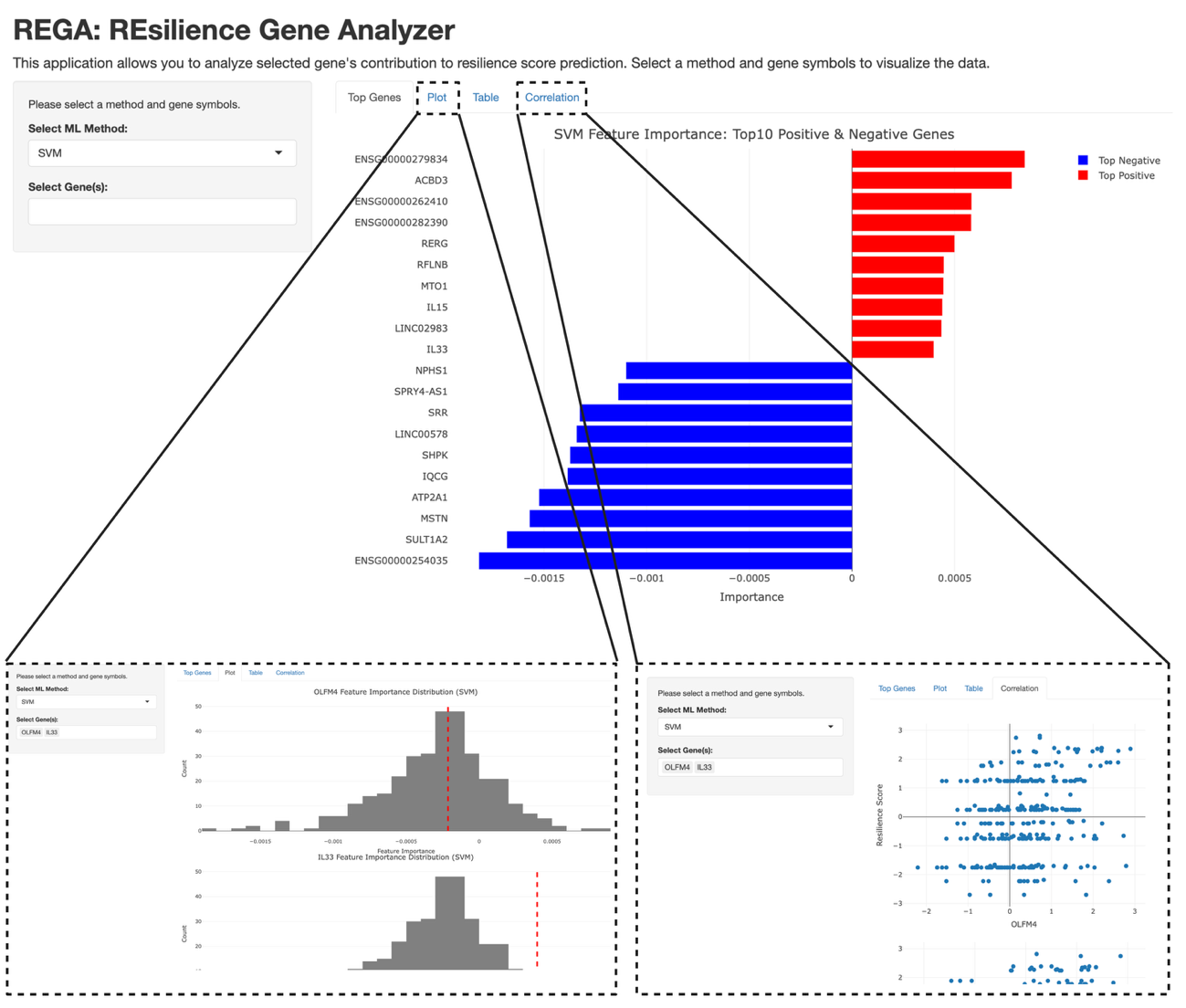

The REGA interface enables visualization of individual gene contributions to CR prediction. Users can select a machine learning model, including ElasticNet, SVM, Random Forest, or XGBoost, and genes of interest for analysis. The Top Genes tab displays genes with the strongest positive and negative contributions. The Plot tab shows the distribution of model-derived contribution scores and highlights the selected gene’s position. The Correlation tab presents the relationship between the selected gene and the resilience score.
